# Building compositional tasks with shared neural subspaces

**DOI:** 10.1101/2024.01.31.578263

**Authors:** Sina Tafazoli, Flora M. Bouchacourt, Adel Ardalan, Nikola T. Markov, Motoaki Uchimura, Marcelo G. Mattar, Nathaniel D. Daw, Timothy J. Buschman

## Abstract

Cognition is remarkably flexible; we are able to rapidly learn and perform many different tasks^1^. Theoretical modeling has shown artificial neural networks trained to perform multiple tasks will re-use representations^2^ and computational components^3^ across tasks. By composing tasks from these sub-components, an agent can flexibly switch between tasks and rapidly learn new tasks^4^. Yet, whether such compositionality is found in the brain is unknown. Here, we show the same subspaces of neural activity represent task-relevant information across multiple tasks, with each task compositionally combining these subspaces in a task-specific manner. We trained monkeys to switch between three compositionally related tasks. Neural recordings found task-relevant information about stimulus features and motor actions were represented in subspaces of neural activity that were shared across tasks. When monkeys performed a task, neural representations in the relevant shared sensory subspace were transformed to the relevant shared motor subspace. Subspaces were flexibly engaged as monkeys discovered the task in effect; their internal belief about the current task predicted the strength of representations in task-relevant subspaces. In sum, our findings suggest that the brain can flexibly perform multiple tasks by compositionally combining task-relevant neural representations across tasks.

## Introduction

Humans, and other animals, can combine simple behaviors to create more complex behaviors^5–9^. For example, once we learn to discriminate whether a piece of fruit is ripe, we can reuse this ability as a component of a variety of foraging, cooking, and eating tasks. The ability to compositionally combine behaviors is thought to be central to generalized intelligence in humans^10^ and a necessary component for artificial neural networks to achieve human-level intelligence^11–15^. When artificial neural networks are trained to perform multiple tasks, they reuse representations and computational components in different tasks^2–4^. Whether the brain similarly reuses sensory, cognitive, and/or motor representations across tasks, and how these representations may be flexibly combined, remains unknown. To test this hypothesis, we trained monkeys to perform three different compositionally related tasks.

## Monkeys perform three compositionally related tasks

All three tasks followed the same general structure: animals were presented with a visual stimulus and had to indicate its category with an eye movement (Fig. 1a). The stimuli were parametric morphs, independently varying in both shape and color (Fig. 1b, see Methods for details). Animals performed three categorization tasks. In the Shape-Axis1 (S1) task, they categorized the shape of the stimulus and then responded on Axis 1: when the shape was more similar to a ‘bunny’, the animal made a saccade to upper-left (UL) target, and when the shape was more similar to a ‘tee’, the animal saccaded to the lower-right (LR) target (Fig. 1c, top row). The Color-Axis2 (C2) task required the animal to categorize the color of the stimulus and respond on Axis 2: if the color was ‘red’, they saccaded to upper-right (UR), and if ‘green’, they saccaded to lower-left (LL; Fig. 1c, bottom row). Finally, in the Color-Axis1 (C1) task, the animals categorized the color of the stimulus (as in the C2 task) and responded on Axis 1 (as in the S1 task; red=LR, green=UL, Fig. 1c, middle row). In this way, the tasks were compositionally related – C1 can be considered as combining the color categorization sub-task of C2 with the motor response sub-task of S1.

**Fig. 1.**
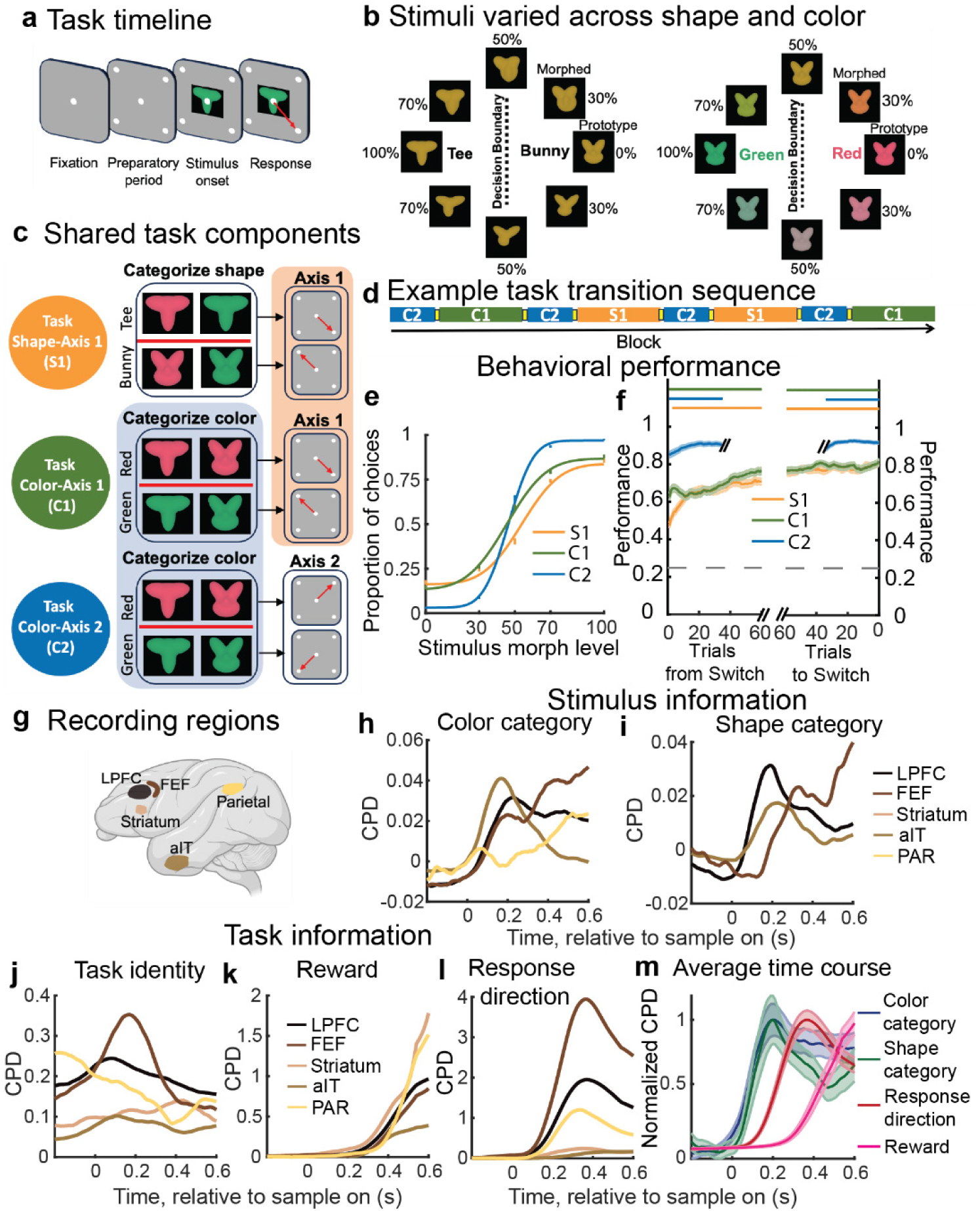
| Monkeys perform three compositional tasks. **a**, Time course of trial. After a brief fixation (500-800ms), a visual stimulus and four response targets appeared on the screen. Animals reported the category of the stimulus with a saccade to one of the four target locations. **b**, Stimuli were morphs in a two-dimensional feature space that varied in shape (left) and color (right). Stimulus categories are indicated by vertical lines and labels. 50% morph stimuli were randomly rewarded. **c**, Schematic of the task design: Task S1 required categorizing the stimulus based on its shape and responding on Axis 1. Task C2 required categorizing the stimulus based on its color and responding on Axis 2. Task C1 required categorizing the stimulus based on its color and responding on Axis 1. Colored backgrounds indicate shared sub-tasks: color categorization sub-task between the C1 and C2 tasks (blue) and response sub-task between S1 and C1 tasks (orange). **d**, Example of task transition sequence. Task switches occurred when performance was equal or greater than 70%. Animals were not instructed as to the identity of the upcoming task, although the axis of response switched between blocks. Number of included blocks for S1/C1/C2 were 94/97/189, respectively. **e**, Average psychometric curve for both animals for the 102/102/51 trials before the switch for S1/C1/C2 tasks, respectively. Number of included blocks for S1/C1/C2 were 94/97/189, respectively. **f**, Average performance of both animals after (left) and before (right) a switch (average window of 15 trials, horizontal line for p ≤ 0.001; binomial test uncorrected for multiple comparisons across trials) **g**, Schematic of locations of neural recordings. **h-l**, To quantify information about task variables in individual neurons, we computed the proportion of variance in neural response uniquely explained by each task variable using the coefficient of partial determination (CPD, average CPD for permuted dataset was subtracted from observed CPD for each neuron, see Methods). Panels show CPD of **h**, color category, **i**, shape category, **j**, task identity, **k**, reward and **l**, response direction. Lines show mean CPDs across all neurons for each region. Regions were omitted if they did not have a significant number of neurons encoding a cognitive variable. **m**, Time course of average normalized CPD across all recorded neurons, normalized by maximum value to show temporal order.

Overall, both animals performed all three tasks well (Figs. 1e and S1; Monkey Si/Ch: S1:81%/77%, C1: 83%/78%, C2: 92%/92%; all p<0.001, binomial test, see Fig. S1 for individual animal performance). Animals performed the same task for a block of trials (Fig. 1d). When they reached a behavioral criterion (performance ≥70%), the task would change (see Methods for details). The animal was not instructed as to the identity of the new task, although the axis of response always changed between blocks (see Methods). So, animals had to learn which task was in effect on each new block. This was reflected in the animal’s behavior, which improved over the first 75 trials of the S1 and C1 tasks (Fig. 1f, left, performance on S1/C1 increased from 0.47/0.62 at trial 15 to 0.71/0.77 at trial 75, both p<0.001, Chi-squared test). As only the C2 task used Axis 2 responses, the animal identified the task quickly, performing it above chance within 15 trials after the switch (Fig. 1f, performance at 15 trials=0.85, p<0.001, binomial test). Altogether, the animals’ behavior suggests they rapidly identified the change in response axis but slowly integrated feedback to learn whether S1 or C1 was in effect^16^.

## Task variables were represented in independent subspaces of neural activity

To understand the neural representations used during each task, we simultaneously recorded neural activity from five cortical and subcortical regions (Fig 1g): lateral prefrontal cortex (LPFC; 480 neurons), frontal eye fields (FEF; 149 neurons), parietal cortex (PAR; 64 neurons), anterior inferior temporal cortex (aIT; 239 neurons) and striatum (caudate nucleus; 149 neurons). All five regions represented task-relevant cognitive variables^17,18^, including the identity of the current task, the color and shape of the stimulus, the response direction, and whether a reward was received (Fig. 1h-1l). The identity of the task was consistently represented throughout the trial (Figs. 1j and S2). After stimulus onset, information about the color (Fig. 1h) and shape of the stimulus (Fig. 1i) was followed by information about the direction of the animal’s response (Fig. 1l), and then the reward received (Fig. 1k, see Fig. 1m for average time course across all regions).

To understand how each task variable was represented in the neural population, we trained classifiers to decode the stimulus color, shape, and motor response from the pattern of neural activity across all neurons (using a pseudo-population to combine neurons from all recording sessions, see Methods for details). Classifiers trained on LPFC neural activity accurately decoded the category of the stimulus’ color in the C1 and C2 tasks (Fig.2a; 79ms, 87ms after stimulus onset for C1 and C2, respectively; similar results were seen when controlling for motor response, Fig. S3a, p<0.001 permutation test). In this way, the classifier defined a “subspace” within the high-dimensional space of neural activity that represented the color category of the stimulus input. Similarly, classifiers trained on LPFC activity decoded the category of the stimulus’ shape during the S1 task (Fig. 2b; 100ms after stimulus onset, p<0.001 permutation test).

**Fig. 2.**
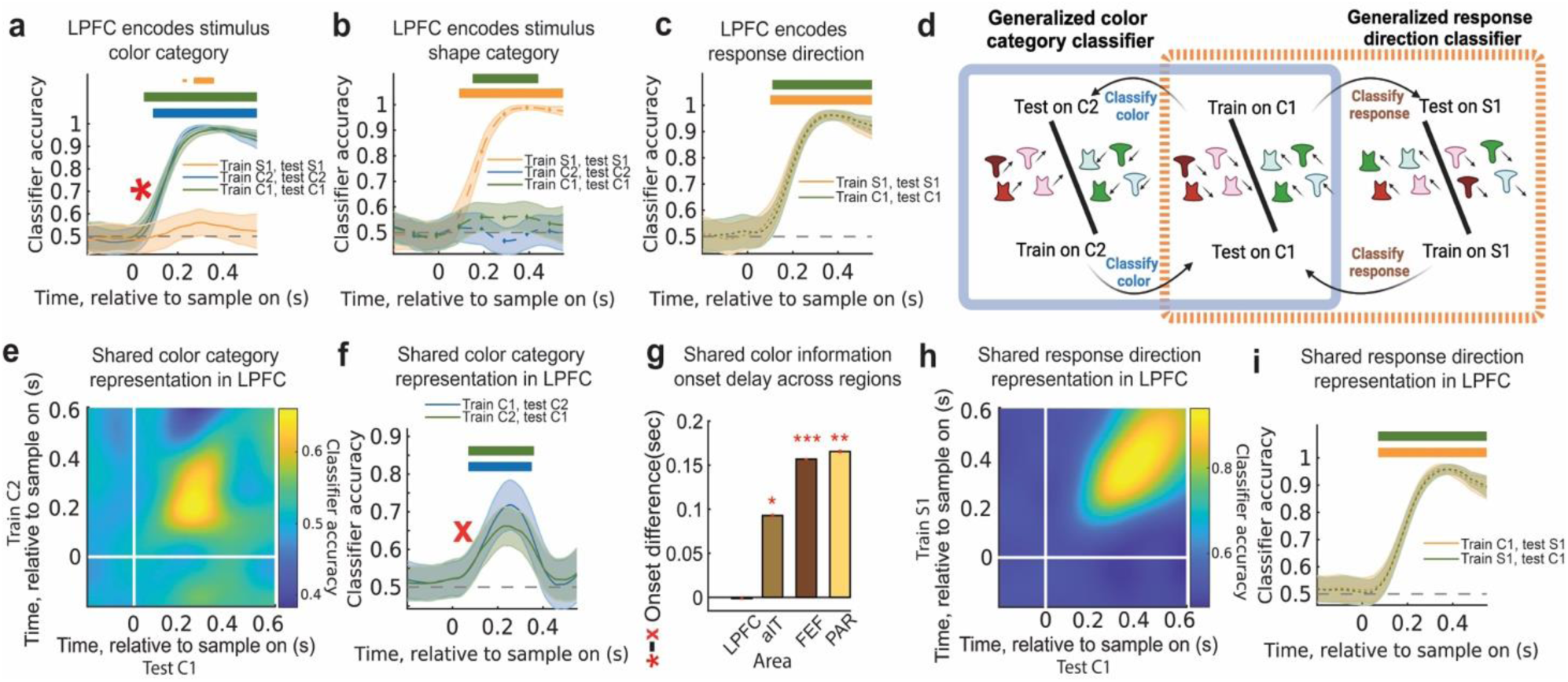
| LPFC encodes shared color and response representations. **a-c**, Accuracy of classifier trained to decode **a**, color category, **b**, shape category, and **c**, motor response from LPFC neural activity. Lines and shading show mean ± s.e.m. over time. Color of line indicates which task was used to train/test the classifier. Distribution reflects 250 resampled classifiers (see Methods for details). Horizontal bars (top right of each plot) indicate above-chance classification (p ≤ 0.05, 0.01, and 0.001 for thin, medium, and thick lines, respectively; permutation test with cluster mass correction for multiple comparisons). **d**, Schematic of classifiers used to test whether color category and response location information were shared across tasks (see Methods). **e**, Cross-temporal cross-task classification accuracy in decoding shared color category in LPFC, trained on C2 task and tested on C1 task. **f**, Time course of cross-task classification accuracy when trained on LPFC neural activity to decode color during C1 task (blue) or C2 task (green) and then tested on the other task. **g**, Difference in the onset of color information during C1 task when decoded from classifiers trained on C1 task versus generalized from C2 task (red star and red x in panels a and f, respectively. p ≤ 0.05, 0.01 and 0.001 for *, **, ***, respectively; t-test). To compare timing, classifier accuracy for each area was smoothed with a 50ms boxcar filter. **h**, Cross-temporal cross-task classification accuracy in decoding shared response direction in LPFC. Trained on S1 task and tested on C1 task. **i**, Time course of cross-task classification accuracy when trained on LPFC neural activity to decode response during C1 task (orange) or S1 task (green) and then tested on the other task.

Task irrelevant stimulus information was attenuated. In LPFC, classification accuracy about the shape of the stimulus was reduced during the C1 and C2 tasks (Fig. 2b) and information about the color of the stimulus was reduced during the S1 task (Fig. 2a). Similarly, task-relevant information, but not task-irrelevant information, was represented in other regions, including in higher-order visual cortex (Fig. S3b-c).

The direction of the animal’s response could be decoded within each task’s response axis (Fig. 2c, see Methods for details). In LPFC, information about the response occurred after the stimulus’ category (115ms, 133ms after stimulus onset in S1 and C1 tasks, respectively; both p<0.001, permutation test, see Fig. S4c for C2 task). Similar results were seen in other regions (Fig. S4a). Altogether, these results show the stimulus color category, stimulus shape category and response direction are broadly represented at the population level in multiple recorded regions.

## Neural representations were shared across tasks

To test whether representational subspaces were reused across tasks, we quantified how well a classifier trained to decode the stimulus’ color category or the animals’ motor response in one task generalized to other tasks (Fig. 2d, Methods). Consistent with a shared representation in LPFC, a classifier trained to decode the color category of a stimulus during the C2 task was able to significantly decode the stimulus’ color category during the C1 task (Figs. 2e and 2f, 65ms after stimulus onset in LPFC, p<0.001, permutation test). The reverse was also true: a classifier trained on C1 could decode stimulus’s color category during the C2 task (Fig. 2f and S5c, 84ms after stimulus onset, p<0.001, permutation test). Importantly, the two tasks require different motor responses and so the shared representation reflected the color category of the stimulus and not the motor response (see Figs. S5a and S6a for further controls for movement). Although color information was reduced in the S1 task, the weak color category representation that did exist also generalized between the C1 and S1 tasks (Fig. S5b).

While color category information was represented in a shared subspace in LPFC, it was represented in task-specific subspaces in other brain regions (Fig. S7). Generalization was weaker in FEF, PAR and IT and was delayed with respect to task-specific sensory information (and was delayed relative to LPFC, Fig. 2g). There was no significant generalization in STR (although stimulus color category could be decoded from those regions, Fig S3b).

Motor response representations also generalized across tasks. A classifier trained to decode response direction in the C1 task generalized to decode response direction in the S1 task (and vice versa; Fig. 2h and 2i, 128/128ms after stimulus onset in LPFC, when training on S1/C1, testing on C1/S1, respectively, p<0.001, permutation test). Unlike stimulus information, the motor representation was shared in all regions (Fig. S4b; perhaps reflecting a widely broadcasted motor signal^19^, but see^20^).

## Shared representations were sequentially engaged during a task

So far, our results suggest the representation of both sensory inputs and motor responses were shared across tasks. In this framework, performing a task requires selectively transforming representations from one subspace to another to support each task^21^. Consistent with this, there was a sequential representation of the stimulus color in the shared color subspace followed by the response in the shared motor subspace during the C1 task (Fig. 3a, 63ms difference in onset time of color response using C2 classifier and motor response using S1, p<0.001 t-test).

**Figure 3.**
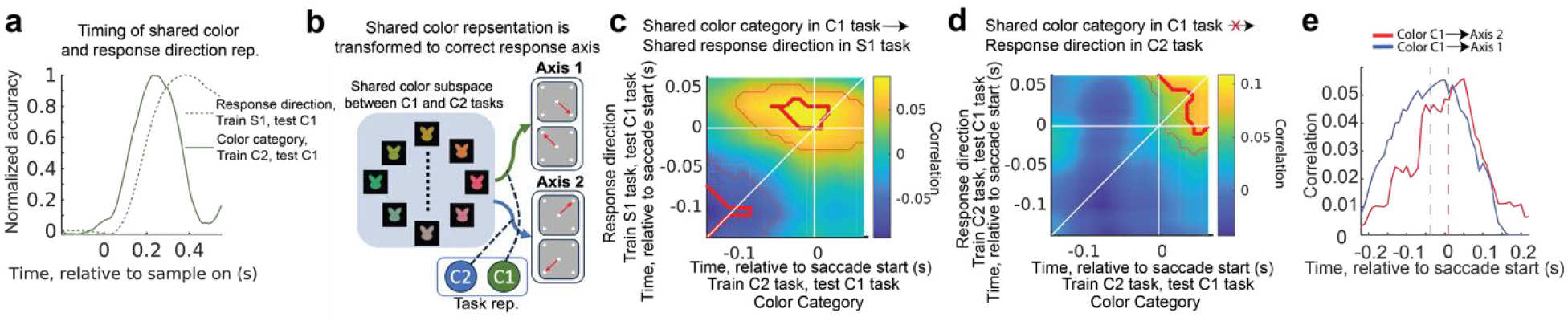
| Shared representations were sequentially transformed during the task. **a**, Sequential processing of shared color category information and shared response direction information in LPFC. Classifier accuracy was normalized to range between maximum accuracy and baseline 200ms before stimulus. **b**, Schematic showing prediction that shared color representation is transformed to the Axis 1 and Axis 2 response axis during the C1 and C2 tasks, respectively. **c**, Cross-temporal, trial-by-trial correlation of shared color category encoding (trained on C2 task, tested on C1 task) and shared response location encoding (trained on S1 task, tested on C1 task). Thin and thick red lines indicate p≤0.05 and p≤0.001, respectively, uncorrected t-test. **d**, Same as panel b, but correlating shared color category encoding (trained on C2 task, tested on C1 task) and response direction encoding on Axis 2 (trained on C2 task, tested on C1 task). **e**, Average cross-temporal correlation along anti-diagonal axis in panel **c** (shared color encoding in C1 task predicts response direction in S1 task, blue line) and panel **d** (shared color encoding in C1 task predicts response direction in C2 task, red line). The shift of the curve’s center towards negative time values reflects the extent to which encoding in the sharedcolor subspace predicts future encoding in the shared response subspace. Dotted lines show mean of gaussian fit to each curve.

To directly test whether this reflected the transformation of information between subspaces, we tested whether information about the stimulus color in the shared color subspace predicted the response in the shared response subspace on a trial-by-trial basis (Fig. 3b).

Figure 3c shows the correlation between the representation of the stimulus’ color category in the C1 task, decoded using the classifier from the C2 task, and the representation of the motor response in the C1 task, decoded using the response classifier from the S1 task (see Methods for details). Correlation was measured across all possible pairs of timepoints, quantifying whether color or motor response representations at one time point were correlated with representations at a future time point. The correlation was shifted upward with respect to diagonal line (36ms before saccade start, Figs. 3c and 3e, blue line), indicating that the encoding in the shared color subspace predicts the future encoding in the shared response subspace.

Importantly, this transformation was specific to the task: during the C1 task the shared color representation was not predictive of the associated response direction along Axis 2 (8ms after saccade start, Figs. 3d and 3e, red line). This is consistent with the shared color representation being selectively transformed into a motor response along Axis 1, and not Axis 2, when the animal was performing the C1 task. In contrast, when the animals performed the C2 task, the shared color representation was transformed into a motor response along Axis 2, and not Axis 1 (Fig. S6b-d).

Together, these results suggest tasks sequentially engage shared subspaces, selectively transforming stimulus representations into motor representations in a task-specific manner. Interestingly, there was a negative correlation between the sensory and motor responses early in the trial (Fig. 3c), which may reflect suppression of the motor response during fixation or integration of the stimulus input.

## Shared subspaces are dynamically engaged by task belief

Theoretical modeling suggests shared subspaces could facilitate cognitive flexibility by allowing the brain to engage previously learned, task appropriate, representational and computational subspaces^2,3^. If true, then this predicts task-appropriate shared subspaces should be engaged as the animal discovers the task in effect.

To begin to test this hypothesis, we first measured the animal’s internal representation of the task (i.e., their ‘belief’ about the task encoded by the neural population). To do so, we trained a classifier to decode the identity of the task using neural activity in LPFC during the fixation period (i.e., before stimulus onset,^22^). Training was restricted to the last 75 trials of the S1 and C1 tasks, when behavioral performance was high (Fig. 1f). We then applied the task classifier to the beginning of the C1 task blocks to measure the animals’ internal belief about the task as they learned which task was in effect (Fig. 4b-1, see Methods for details).

**Fig. 4.**
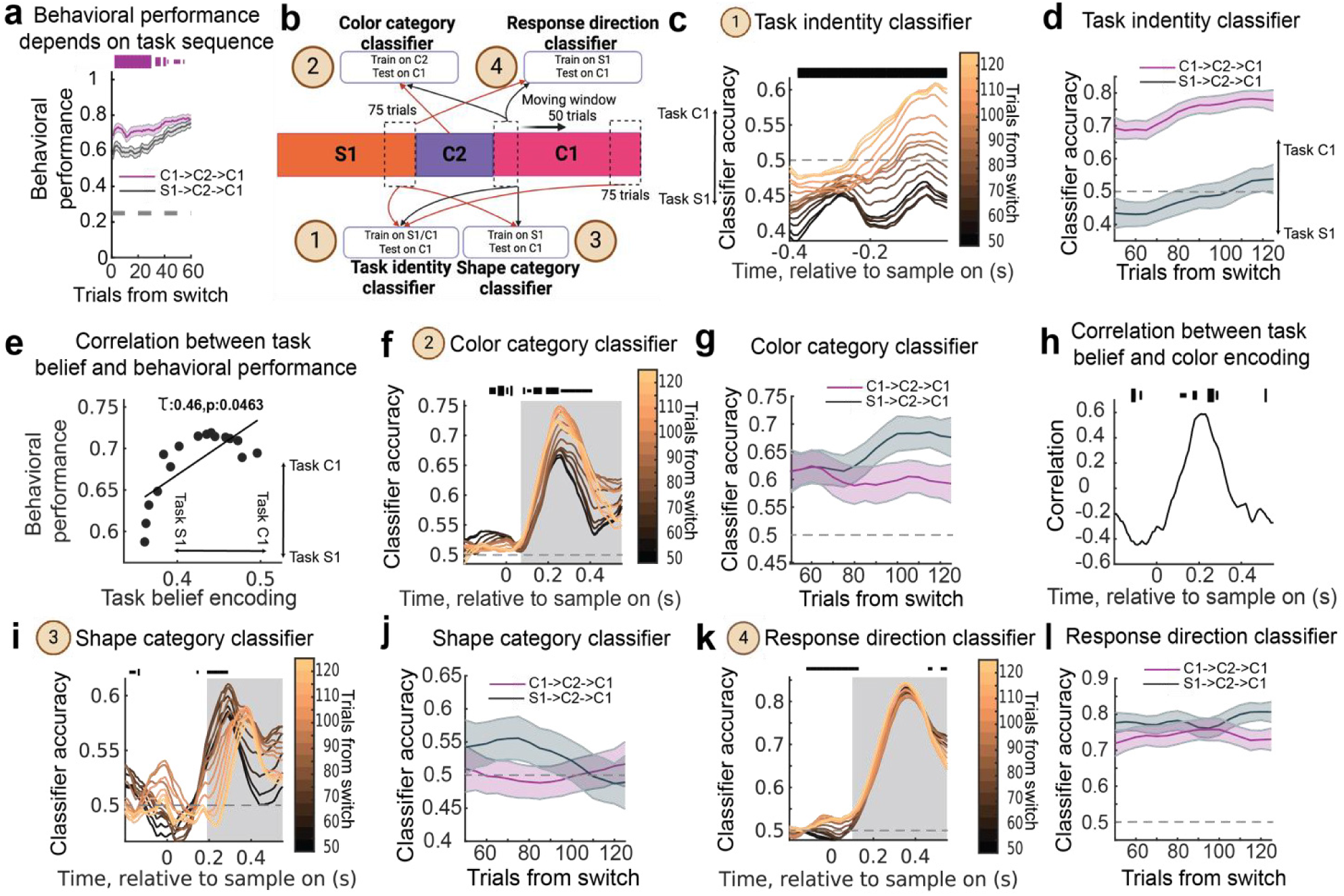
| Shared sub-task subspaces are dynamically engaged during task discovery. **a**, Behavioral performance of monkeys during the C1 task depended on the sequence of preceding tasks. Dashed gray line is 25% chance level. Horizontal bars indicate significant difference between performance in two sequences (p ≤ 0.05, 0.01, and 0.001 for thin, medium, and thick lines, respectively; Chi-squared test uncorrected for multiple comparisons across trials). Number of included blocks for C1-C2-C1/S1-C2-C1 sequences were 44/53, respectively. **b**, Schematic of classifier tests during task discovery.1: Task identity classifier was trained to decode task identity on 75 trials before switch in S1 task and C1 task, tested on sliding window of 50 trials starting from switch into C1 task. 2: Color category classifier was trained to decode color category on 75 trials before switch in C2 task, tested on sliding window of 50 trials starting from switch into C1 task. 3: Shape category classifier was trained to decode shape category on 75 trials before switch in S1 task, tested on sliding window of 50 trials starting from switch into C1 task. 4: Response direction classifier was trained to decode response direction on 75 trials before switch in S1 task, tested on sliding window of 50 trials starting from switch into C1 task. Same test trials were used for 1, 2 and 3 classifiers. Red and black arrows denote train and test trials, respectively. **c**, Classifier accuracy for task identity during –400ms to 0ms period from sample onset. Darker to lighter color show progression in trial blocks of 50 shifted by 5 trials from switch trial in S1-C2-C1 task transition. Lines show mean classification accuracy after stimulus onset for 250 iterations of classifiers. Horizontal bars indicate above chance trend for classifier accuracy during task discovery (p ≤ 0.05, 0.01, and 0.001 for thin, medium, and thick lines, respectively; modified Mann-Kendall test, Benjamini-Hochberg correction for multiple comparisons across time) **d**, Comparison of progression of task identity classifier accuracy in C1-C2-C1 task sequence (purple) and S1-C2-C1 task sequence (black). Average accuracy was computed using classifier performance in –400ms to 0ms before stimulus onset period. Lines and shading show mean ± s.e.m. classification accuracy after stimulus onset. Distribution reflects 250 iterations of classifiers. **e**, Correlation of behavioral performance and task belief encoding. **f**, As in panel c, but showing classifier accuracy for color category classifier in S1-C2-C1 task transition. Grey box shows period with significant color category information (see Methods). **g**, Comparison of progression of color category classifier accuracy in C1-C2-C1 sequences (purple) and S1-C2-C1 sequences (black). Average accuracy was computed using classifier accuracy in 100ms to 300ms from stimulus onset. **h**, Correlation between task belief encoding and color category encoding (see Methods for details). p ≤ 0.05, 0.01, and 0.001 for thin, medium, and thick lines, modified Mann-Kendall test, Benjamini-Hochberg correction for multiple comparisons across time. **i,j** same as **f,g** but for shape category classifier. Grey box shows period with significant shape category information. **k,l** same as **f,g** but for response direction classifier. Classifier accuracy was computed using classifier accuracy in 200ms to 400ms after stimulus onset. Grey box shows period with significant response direction information.

As noted above, monkeys slowly learned whether the S1 or C1 task was in effect^16^. Interestingly, animals’ rate of learning depended on the sequence of tasks. When switching into a C1 task, animals learned more quickly when the previous Axis 1 task was C1 compared to S1 (Fig. 4a, purple: C1-C2-C1 task sequence, black: S1-C2-C1; Δ=10.26% change in performance over first 20 trials, p<0.001, Chi-squared test). A similar pattern was seen for S1: performance on task sequence S1-C2-S1 was greater than C1-C2-S1 (Fig. S8a, Δ=4.99%, p=0.032, Chi-squared test). As the C2 task had a unique response axis, it was not affected by the preceding task (Fig. S8b Δ=1.42% between S1-C2 and C1-C2 sequences, p=0.234, Chi-squared test).

As with behavior, the performance of the task classifier increased over trials (Fig. 4c) and depended on the sequence of tasks (Fig. 4d). During S1-C2-C1 sequences, the classifier was slightly below chance initially, suggesting a slight encoding of the S1 task, before increasing as the animals learned (Fig. 4c-d, from 43% to 54% from trial 50 to 125, p<0.001 modified Mann-Kendall trend test). On C1-C2-C1 task sequences, classifier performance was high immediately after the switch to C1 (69% classifier accuracy on trial 50, p<0.001, permutation test) and slightly increased with trials (Fig. 4d, purple line; 8% increase from trial 50 to 125, p<0.001 modified Mann-Kendall trend test). Moreover, the task belief during the C1 task was correlated with behavioral performance on the color categorization task (Fig. 4e, τ=0.46, p=0.0463 modified Mann-Kendall trend test) and anti-correlated with how much their behavior depended on the shape category (Fig. S9a, τ=-0.55, p=0.02 modified Mann-Kendall trend test). Together, these results suggest the monkeys tracked whether the S1 or C1 task was in effect, and that this belief was at least partially maintained between blocks of the Axis 1 tasks, leading to a bias in the neural representation and behavior.

Given this measure of the animal’s internal representation of the task, we next asked whether this representation predicted the strength of representations in the shared color category and shared motor response subspaces. Similar to the task representation, the strength of the representation in the shared color subspace increased as the animals discovered the C1 task (Fig. 4f, Fig. 4b-2, see Methods for details). Like task representations, color representations depended on task sequence (Fig. 4g). Shared color representations increased during the C1 task when it followed a S1-C2-C1 sequence (62% to 68%, Δ=6% from trial 50 to 125, p<0.001 modified Mann-Kendall trend test), but remained stable during C1-C2-C1 sequences (62% to 59%, Δ=3% from trial 50 to 125, p=0.12 modified Mann-Kendall trend test). This suggests that shared color subspace is dynamically engaged as the animal updates its belief about the current task. Indeed, during learning, the strength of internal task representation in LPFC during fixation was correlated with the strength of shared color category representation after onset of the stimulus (Fig. 4h, see Methods for details).

The animal’s behavior and internal representation of the task suggest they initially expected to perform the S1 task in S1-C2-C1 task sequences. If true, and task representations modulate the strength of shared subspace representations, then shape information should initially be strong and then decay as the animal learns the C1 task is in effect. To test this, we trained a classifier to decode the stimulus’ shape category during the last 75 trials of the S1 task, and then tested it as the animal discovered the C1 task (Fig. 4i, Fig. 4b-3, see Methods for details). In contrast to color, shape representation was significant immediately after the switch to the C1 task during S1-C2-C1 sequences (54%, p=0.02, permutation test) and significantly decreased as the animal learned (Fig. 4i-j, 49%, Δ=5%, p=0.049, modified Mann-Kendall trend test). In contrast, shape information was reduced overall and remained stable during C1-C2-C1 task sequences (Fig. 4j, purple, 51% on trial 50, Δ=1%, p=0.203, modified Mann-Kendall trend test). Finally, consistent with task belief modulating engagement of shape representations, belief about the C1 task was inversely correlated with the representation of shape (Fig. S9b).

In contrast to shared color and shape subspaces, the animals’ motor response was stably decoded in a shared subspace (Fig. 4k-l, Fig. 4b-4, see Methods for details, Δ=3%, p=0.106, and Δ=1%, p=0.284, for S1-C2-C1 and C1-C2-C1 sequences, respectively; modified Mann-Kendall trend test; see Methods for details). The representation in the shared response subspace was also not correlated with the strength of internal task representation in LPFC (τ=0.38, p=0.070, modified Mann-Kendall trend test, Fig. S9c).

Altogether, these results suggest the animals use their internal belief to selectively engage the relevant shared color and shape subspaces during the C1 task. The task-relevant stimulus subspace (color) was then dynamically projected onto the shared representation of the response.

## Task belief scaled representations within the shared subspaces

Previous work suggests the gain of stimulus features can change, depending on their relevance for the current task^23–26^. Indeed, we found task-irrelevant information was attenuated (Fig. 2a, 2b). Furthermore, this attenuation depended on the animals’ internal belief about the task: as they discovered the C1 task was in effect, the representation of color category was magnified (Fig. 5a), while shape representation was attenuated (Fig. 5b).

**Fig. 5.**
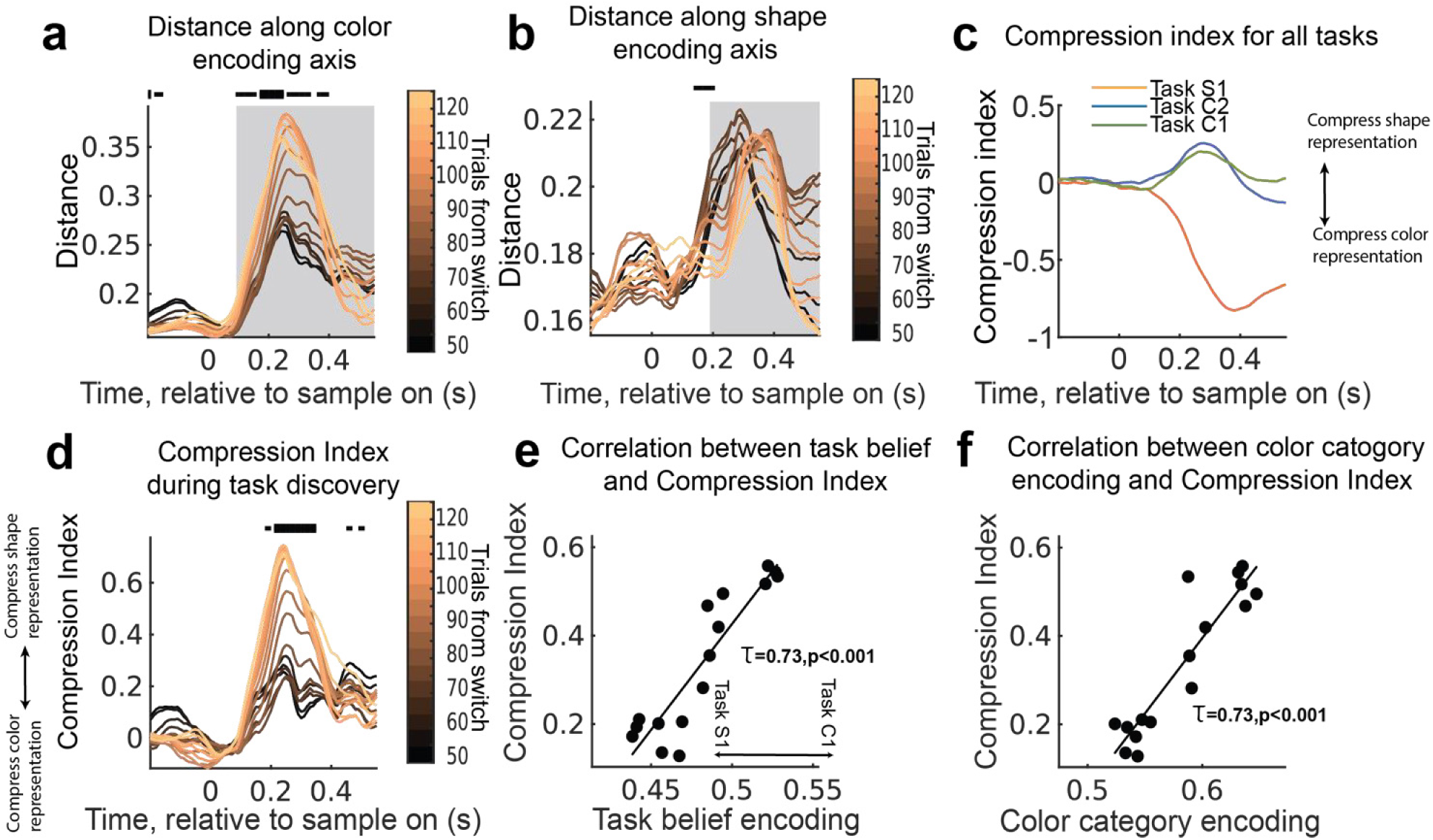
| Stimulus representations were compressed during flexible behavior. **a**, Distance between stimuli along color category encoding axis (Methods). Darker to lighter color show progression in trial blocks of 50 shifted by 5 trials from switch trial in S1-C2-C1 sequences. Lines show mean value for 250 iterations of classifiers. Horizontal bars indicate significant trend in distance during task discovery (p ≤ 0.05, 0.01, and 0.001 for thin, medium, and thick lines, respectively; modified Mann-Kendall test, Benjamini-Hochberg correction for multiple comparisons across time). Grey box shows period with significant color category information. **b**, same as **a** but for distance along shape category encoding axis.Grey box shows period with significant shape category information. **c**, Compression Index during trial for three tasks. Compression index is taken as the log of the ratio of the separability of stimuli in color and shape subspaces. **d**, Compression Index during task discovery in S1-C2-C1 sequences. **e**, Correlation between task belief encoding and average compression index during 100ms-300ms since stimulus onset (Methods). **f**, Correlation between color category encoding and average compression index during 100ms-300ms since stimulus onset (Methods).

Such amplitude modulation can be thought of as a geometric scaling of stimulus representations in feature space^23,24,27^. To quantify this, we defined a ‘Compression Index’ (CPI) as the log of the ratio between the separability of stimuli in (i) the color subspace and (ii) the shape subspace. Color and shape representations were scaled in all three tasks. There was greater separation of color representation during C1 and C2 tasks, and greater separation of shape representations during the S1 task (Fig. 5c). This scaling changed as animals learned which task was in effect (Fig. 5d). In fact, CPI was positively correlated with the strength of the task representation and color category encoding in LPFC (Fig. 5e and 5f, τ=0.73, p<0.001 and τ=0.73, p<0.001, respectively).

Together, these results suggest scaling of stimulus information is a common computation across all the three tasks that can be flexibly engaged based on the monkeys’ belief about the current task, dynamically adjusting the representational geometry of sensory representations^28–30^.

## Discussion

Our results suggest the brain can perform a task by compositionally combining a set of representational subspaces. We found ‘shared’ subspaces of neural activity in prefrontal cortex that represented sensory inputs and motor actions across multiple tasks. This contrasts with previous work that has found high-dimensional, non-linear representations that ‘mix’ sensory, motor, and task information in a unique way for each task^31,32^. Such non-linear, high-dimensional, representations are easy to linearly separate, facilitating learning and performing arbitrary tasks^33^. However, because each task representation is essentially unique, learning cannot be generalized across tasks^24^. Instead, our results are consistent with recent theoretical models that suggest neural representations and computational components are reused across tasks. Shared representational subspaces could speed learning by allowing knowledge to generalize across tasks^4^. For example, the neural representation (and associated computations) that were learned to categorize the color of a stimulus during C2 could be generalized to support behavior during C1. Future work is needed to test this hypothesis as subjects learn new tasks.

While sharing representations across tasks may facilitate generalization between tasks, it could also lead to interference. Given that the animals were trained for months on the task, one might expect the brain to form independent task representations in order to reduce interferences between tasks^34,2^. One potential explanation for our observation that representations are shared across tasks could be the uncertainty in which task to perform (as there was no cue for the current task). This may lead to continual learning, which would be facilitated by shared representations^35^. Instead, we found task-irrelevant dimensions were compressed^28,36^ as a function of the animals’ internal belief about the task state (Fig. 5). Compression could reduce interference from non-relevant stimulus inputs (specifically, incongruent stimuli in the task).

Generalization of subspaces across tasks was strongest in prefrontal cortex. This is consistent with previous work that has found projecting multiple stimuli^37^ or memories^38^ into a common subspace within prefrontal cortex can allow a single task to generalize across stimuli and/or memories^39^. Our results show subspace representations within prefrontal cortex can be shared across multiple tasks. Of course, other regions may also be involved – for example, previous work has found generalization of representations in the hippocampus^37,40^.

Our results are consistent with prefrontal cortex acting as a ‘global workspace’^41^, with different subspaces of neural activity representing different task-relevant information. In this model, performing a task requires transforming information from the relevant sensory subspace to the appropriate motor response subspace. Consistent with this, we found sensory representations in the category subspace predicted responses in the motor subspace on a trial-by-trial basis (Fig. 3). The mechanism by which information is transformed between subspaces remains unknown, although previous work suggests rotation of neural representations may be one possible mechanism^42,43^.

Which subspaces were engaged, and how information was transformed, depended on the animals’ internal belief about the current task (Figs. 3 and 4). This is consistent with computational models of cognitive flexibility that suggest task representations can act as a control input to select task-appropriate representations and computations in a neural network (whether feed-forward^26^ or recurrent^2,3,44^). Thus, if the necessary representations already exist, then learning a new task only requires discovering the appropriate control input – i.e., the representation of the task that engages the appropriate representations/computations. This could dramatically speed learning, either by learning the task representations through reward feedback^45,46^ or recalling it from long-term memory^47^.

## Acknowledgements

We thank Britney Morea, Neeraja Rajagopalan for assistance with monkeys; Srdjan Ostojic, Carlos Brody, Tatiana Engel and Christopher M. Langdon for thoughtful discussions; Tatiana Engel, Caroline Jahn, Qinpu He, Polina Lamshchinina, Iman Wahle, Seth Akers-Campbel and Junchol Park for feedback on the manuscript and the Princeton Laboratory Animal Resources staff for support. This work was supported by NIH R01MH129492 (T.J.B.).

## Author contributions

TJB and ST conceived the project. ST collected experimental data with assistance from NTM, MU and TJB. ST analyzed the data with input from TJB, AA, FMB, MM, and NDD. ST wrote first draft of the paper. ST, TJB, FMB, MM, AA, NDD, NTM and MU edited the paper. TJB acquired funding. TJB supervised the project.

## Supplemental Figures

**Fig. S1.**
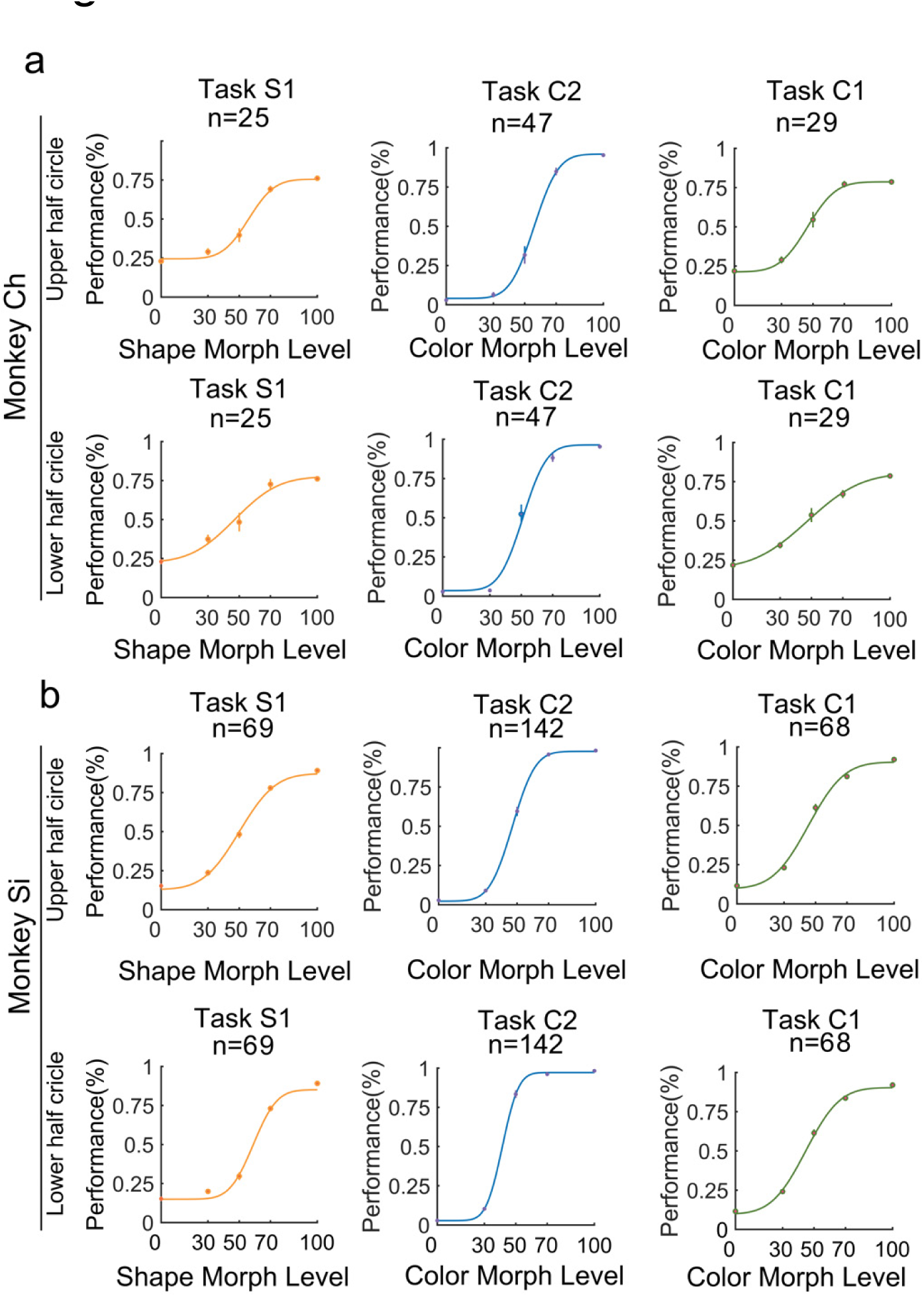
| Both monkeys perform the task well. Follows Fig. 1e. Average psychometric curve for (**a**) monkey Ch and (**b**) monkey Si. Behavioral performance is shown for the last 102 trials of a bock for the S1 and C1 tasks and last 51 trials of blocks of the C2 task. N indicates the number of blocks per task.

**Fig. S2.**
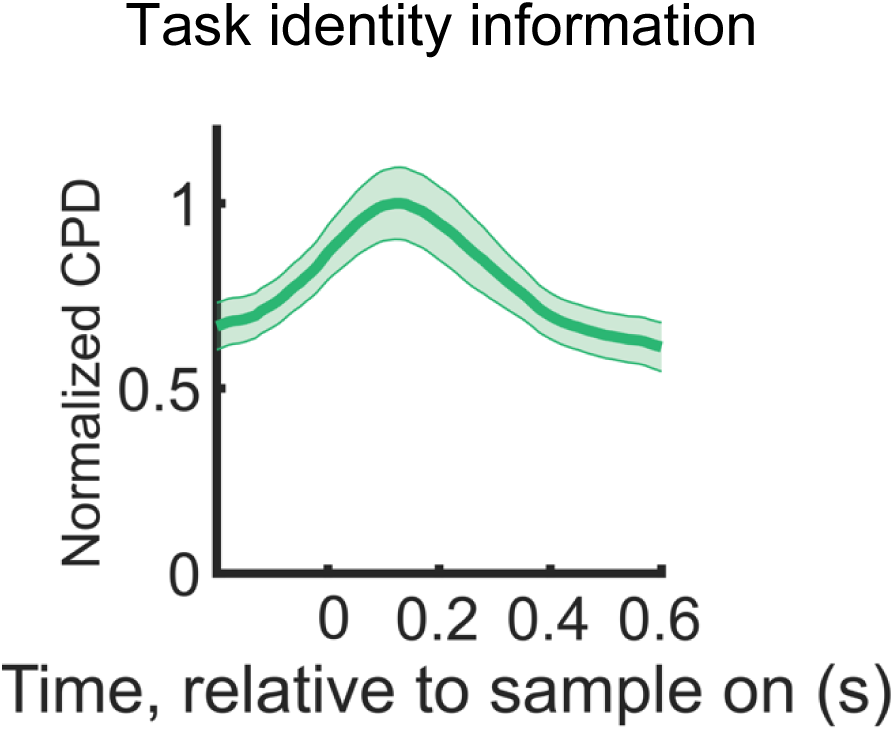
| Task identity is represented throughout the trial. Follows Fig. 1m. Normalized CPD for task identity averaged for all regions. Lines and shading show mean± s.e.m.

**Fig. S3.**
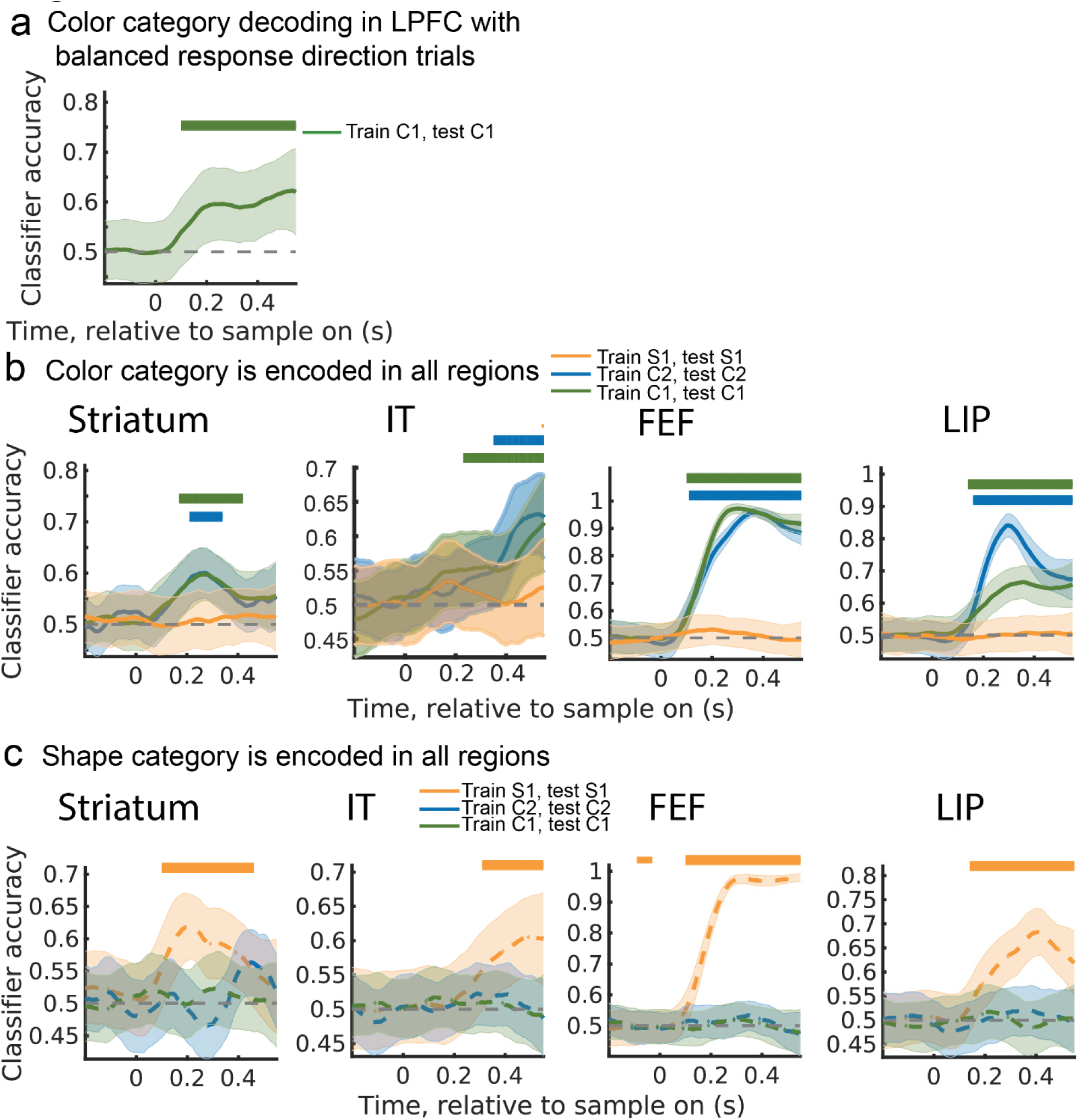
| Color and shape category were decoded in all regions. **a**, Follows Fig. 2a. Time course of accuracy of classifier trained to decode the color category of the stimulus based on neural activity in LPFC. To control for movement, the number of trials with each response direction were balanced within each color category. Only trials with responses on the correct response axis were included (e.g., Axis 1 for task C1). **b**, Follows Fig. 2a. Time course of accuracy of classifier trained to decode the color category of the stimulus. Classifiers were trained for each task and tested on withheld trials of the same task (colored lines). Classifiers were trained separately for each brain region (Striatum, IT, FEF, and LIP in each column, moving left to right). Lines and shading show mean ± s.e.m. classification accuracy after stimulus onset. Distribution reflects 250 iterations of classifiers. Horizontal bars (top right of each plot) indicate above-chance classification (p ≤ 0.05, 0.01, and 0.001 for thin, medium, and thick lines, respectively; permutation test with cluster mass correction for multiple comparisons). **c**, Follows Fig. 2b. Same as **b**, but for shape category information.

**Fig. S4.**
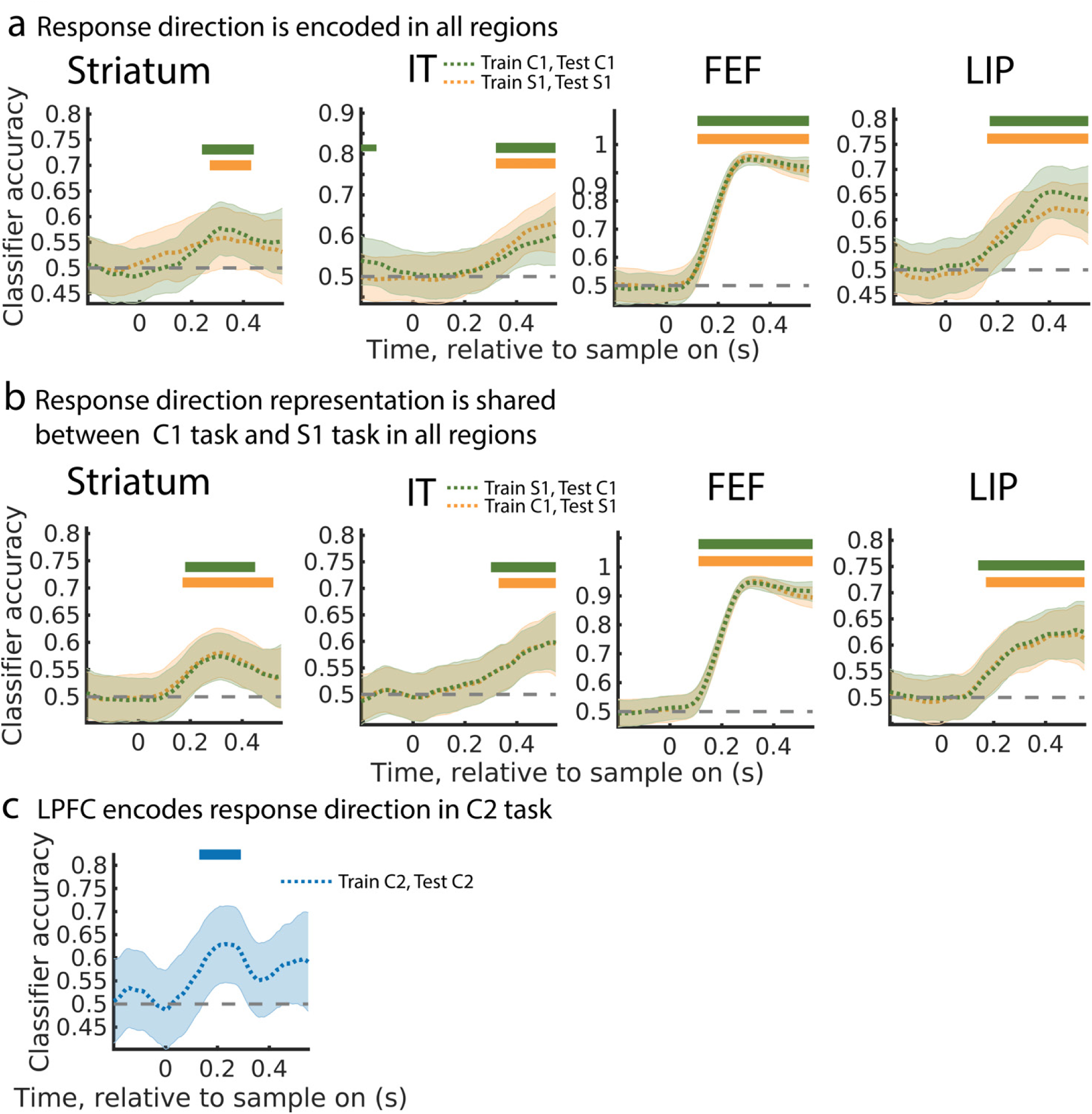
| Response direction was decoded in all regions. **a**, Follows Fig. 2c. Time course of accuracy of classifier trained to decode response direction for S1 and C1 tasks. Classifiers were trained separately for each brain region (Striatum, IT, FEF, and LIP in each column, moving left to right). Lines and shading show mean ± s.e.m. classification accuracy after stimulus onset. Distribution reflects 250 iterations of classifiers. Horizontal bars along top indicate above-chance classification (p ≤ 0.05, 0.01, and 0.001 for thin, medium, and thick lines, respectively; permutation test with cluster mass correction for multiple comparisons). **b**, Follows Fig. 2i. Same as **a**, but showing timecourse of accuracy of classifier trained on S1 task (green) or C1 task (orange) and then tested on the other task (C1 and S1, respectively). **c**, Follows Fig. 2c. As in **a**, but showing the time course of accuracy for a classifier trained to decode response direction during the C2 task from neural activity in LPFC.

**Fig. S5.**
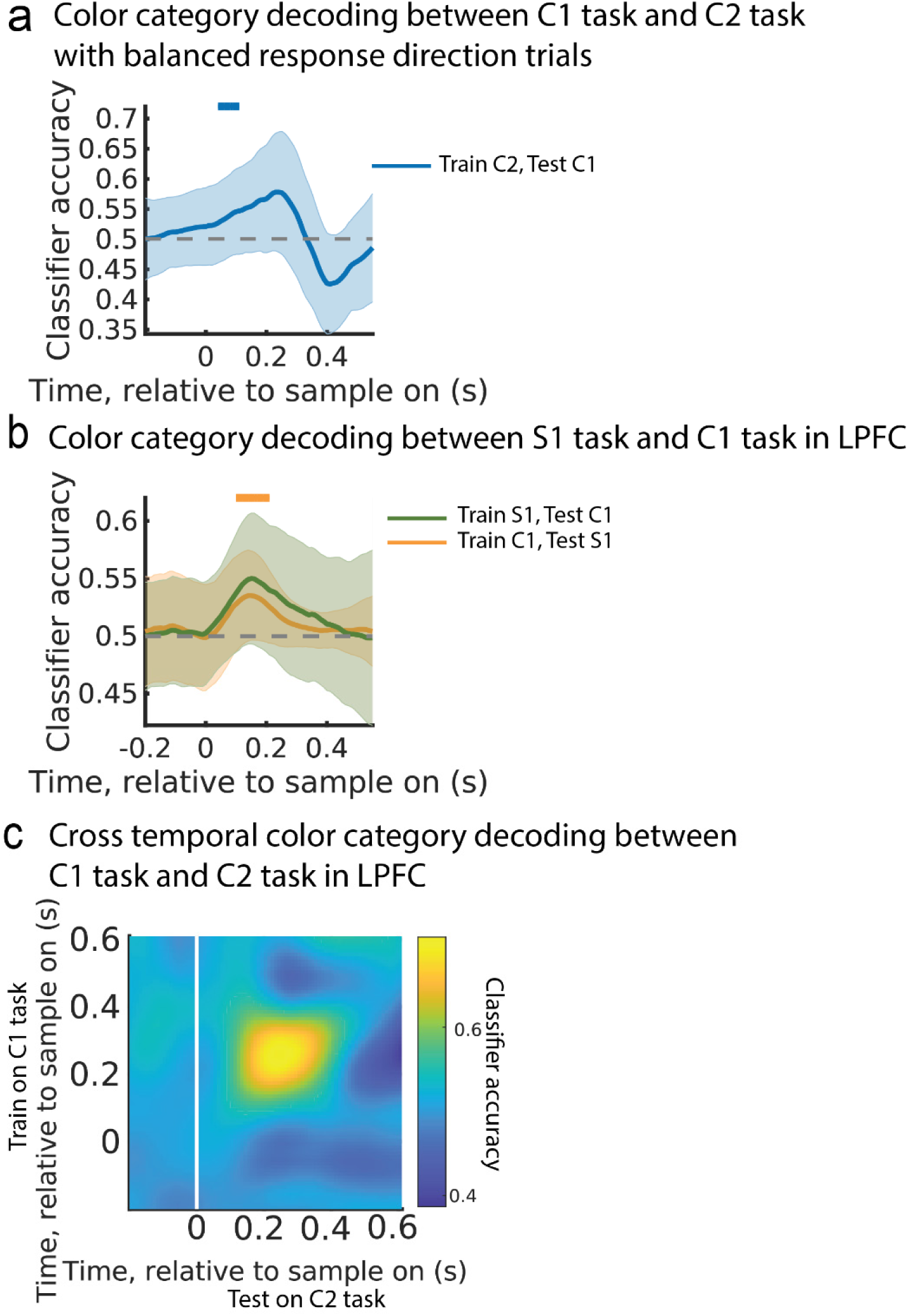
| Color category was cross decoded between tasks in LPFC. **a**, Time course of accuracy of classifier trained to decode color category on trials of the C1 task, while balancing motor response (as in Fig. S3a). Classifier is then tested on the C2 task. Note, reduction in decoding accuracy reflects the reduced number of trials and introduction of error trials. Lines and shading show mean ± s.e.m. classification accuracy after stimulus onset. Distribution reflects 250 iterations of classifiers. Horizontal bars along top indicate above-chance classification (p ≤ 0.05, 0.01, and 0.001 for thin, medium, and thick lines, respectively; permutation test with cluster mass correction for multiple comparisons). **b**, As in Figure 2f, but showing the accuracy of classifiers trained to decode color category from LPFC neural activity during the C1 task (orange) and S1 task (green) and then tested on the other task (S1 and C1, respectively). Classifiers were trained only on correct trials. Plot style follows panel a. **c**, As in Figure 2e, but showing the cross-temporal cross-task classification accuracy in decoding shared color category in LPFC when training on C1 task trials and testing on C2 task trials.

**Fig. S6.**
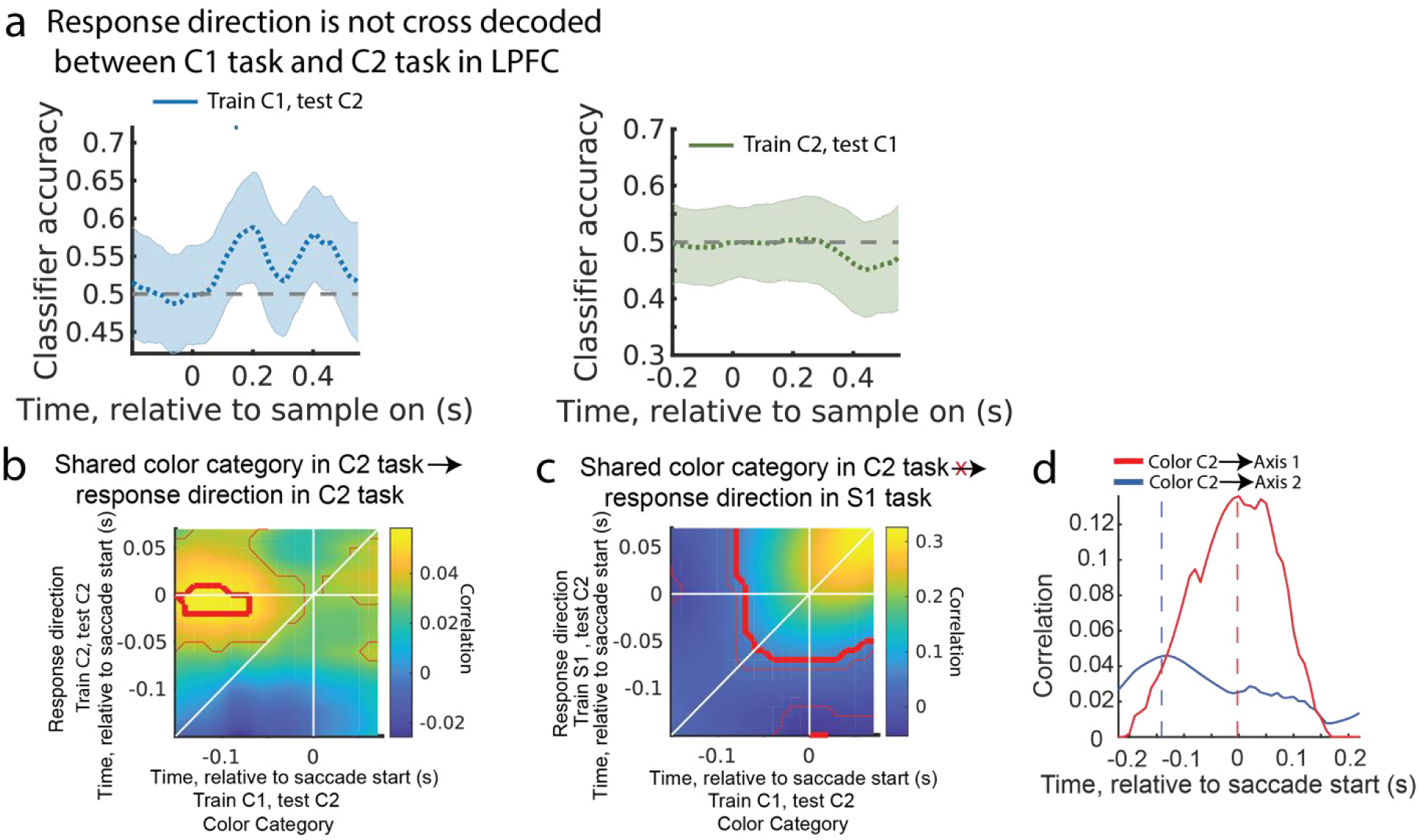
| Response direction is not cross decoded between axes. **a**, Classifiers are unable to decode response direction between axes. Classifiers were trained to decode response direction from LPFC neural activity during the C1 task (left) or C2 task (right) and then tested on accuracy to decode the matched hemifield location during the C2 task (left) or C1 task (right). Lines and shading show mean ± s.e.m. classification accuracy after stimulus onset. Distribution reflects 250 iterations of classifiers. Horizontal bars indicate above-chance classification (p ≤ 0.05, 0.01, and 0.001 for thin, medium, and thick lines, respectively; permutation test with cluster mass correction for multiple comparisons). **b**, Shared color category in C2 task was transformed into response on Axis 2 and not Axis 1. As in Fig. 3c, but shows cross-temporal correlation across trials of shared color category encoding (trained on C1 task, tested on C2 task) and response direction encoding on Axis 2 (trained on C2 task, tested on C2 task). **c**, As in Fig. 3d, but correlating shared color category encoding (trained on C1 task, tested on C2 task) and response direction encoding on Axis 1 (trained on S1 task, tested on C2 task). **d**, As in Fig. 3e, but average cross-temporal correlation along anti-diagonal axis in panel b (shared color encoding in C2 task predicts response direction in C2 task, blue line) and panel c (shared color encoding in C2 task predicts response direction in S1 task, red line). Thin and thick red lines in panels b, c indicate p≤ 0.05 and p≤ 0.001, respectively, uncorrected t-test.

**Fig. S7.**
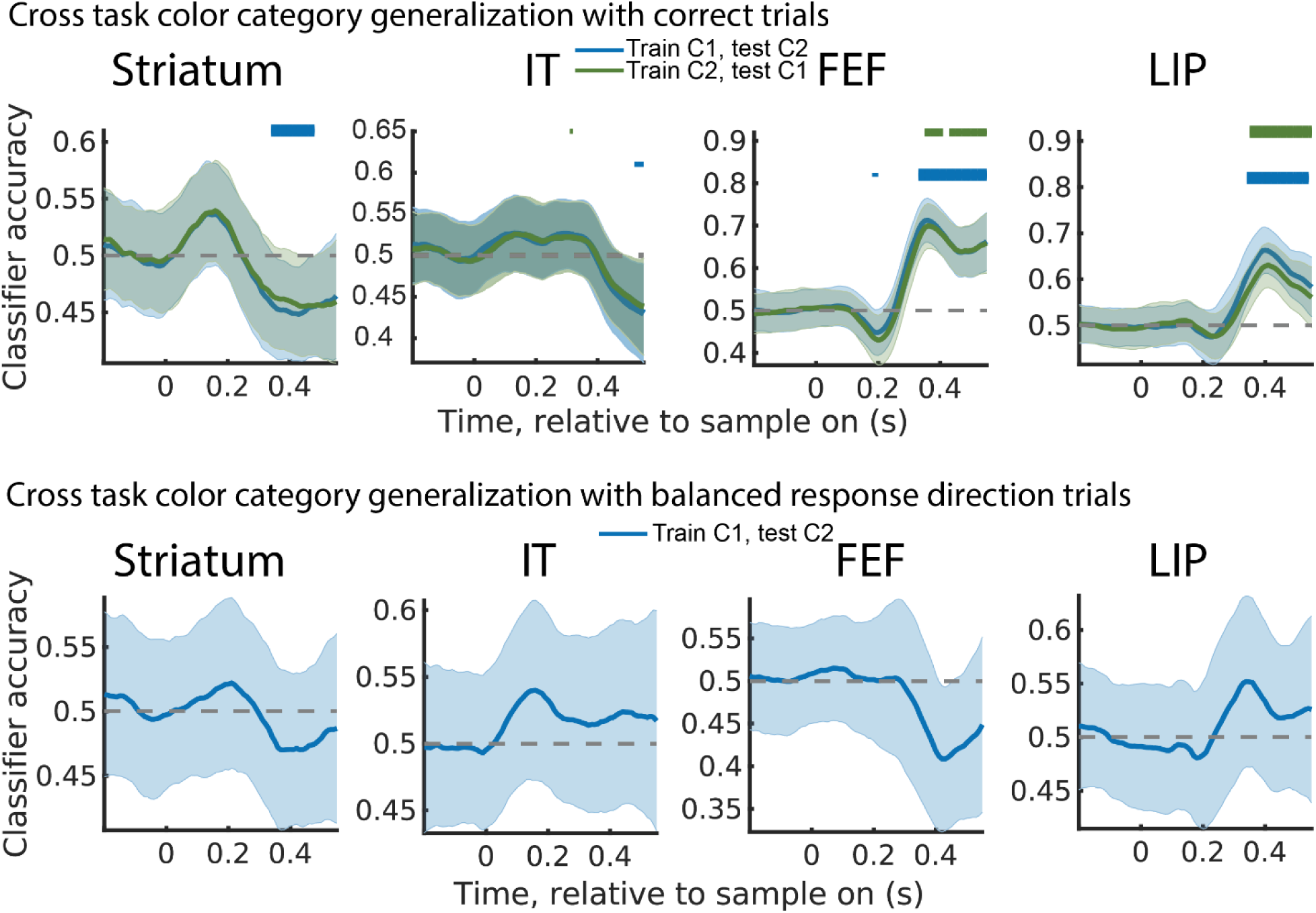
| Color category was shared across tasks in a subset of regions. **top**, Follows Fig. 2f. Time course of accuracy of classifier trained to decode the color category during the C1 task (blue) and C2 task (green), and then tested on the other task (C2 and C1, respectively). Independent classifiers were trained for each brain region: striatum, IT, FEF and PAR (from left to right, across columns). Classifiers were trained on correct trials alone. lines and shading show mean ± s.e.m. classification accuracy after stimulus onset. Distribution reflects 250 iterations of classifiers. Horizontal bars indicate above-chance classification (p ≤ 0.05, 0.01, and 0.001 for thin, medium, and thick lines, respectively; permutation test with cluster mass correction for multiple comparisons). **bottom**, Same as top, but using balanced response direction trials in each color category.

**Fig. S8.**
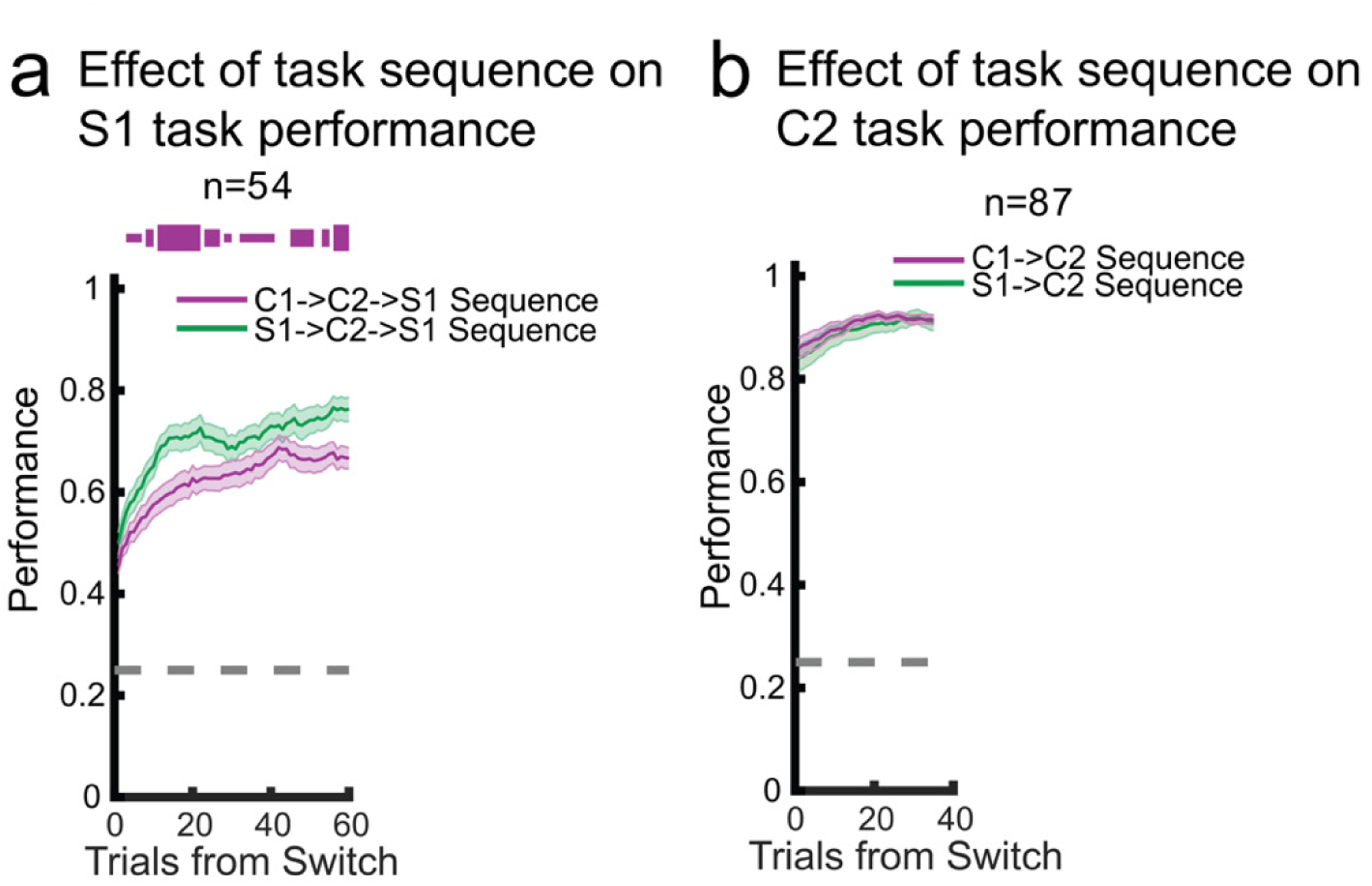
| Task sequence affected behavioral performance in S1 and C1 tasks but not in C2 task. **a**, Follows Fig. 4a. Comparison of average behavioral performance of both monkeys on S1 task when at the end of a sequence of S1-C2-S1 tasks or C1-C2-S1 tasks. Horizontal bars indicate above chance difference between performance in two sequences, p ≤ 0.05, 0.01, and 0.001 for thin, medium, and thick lines, respectively. Chi-squared test uncorrected for multiple comparisons across trials. Number of included blocks for C1-C2-C1/S1-C2-C1 sequences were 40/54, respectively. **b**, Same as **a**, but comparing average behavioral performance of both monkeys for S1-C2 and C1-C2 task sequences. Number of included blocks for C1-C2-C1/S1-C2-C1 sequences were 82/87, respectively.

**Fig. S9.**
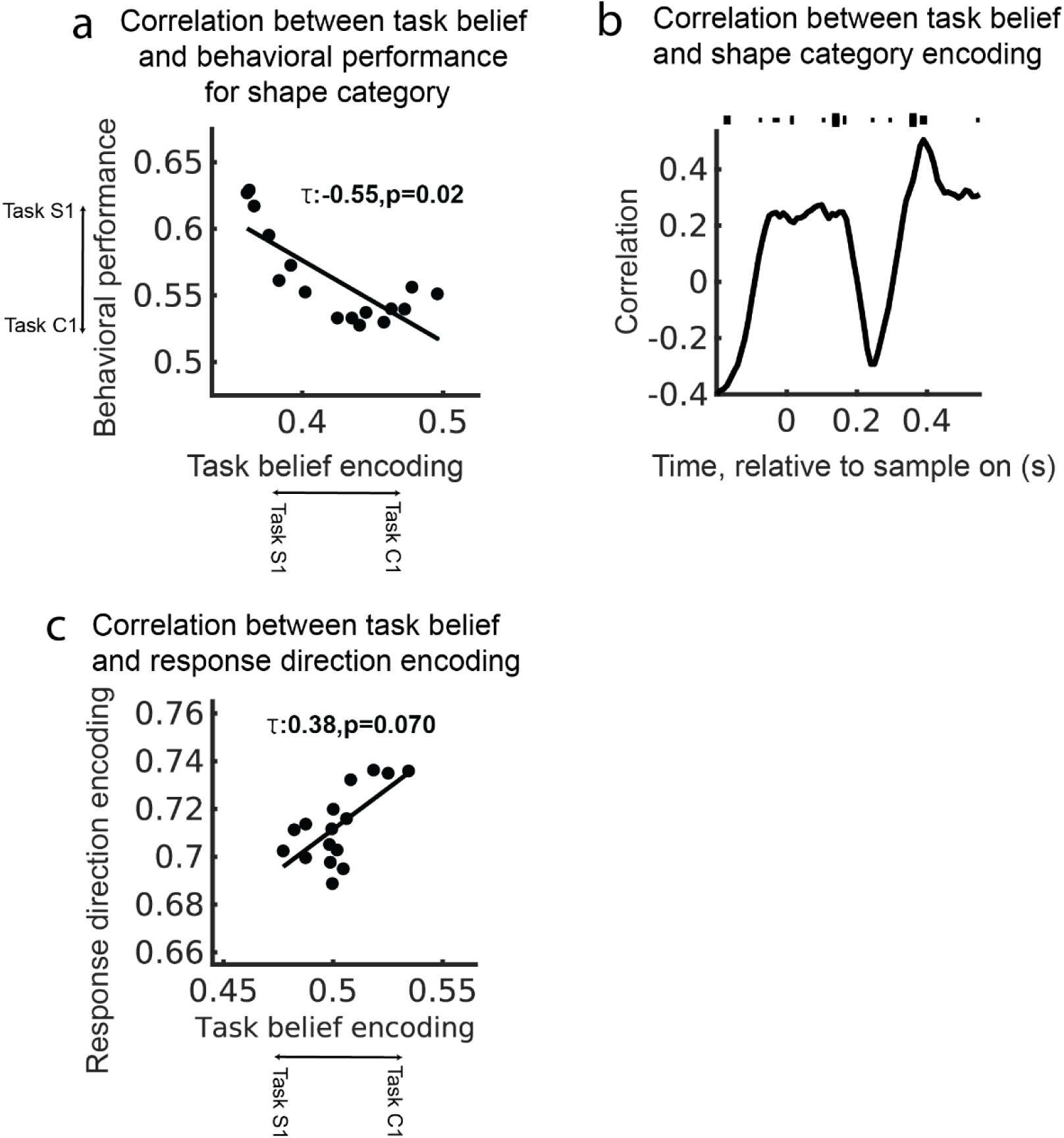
| During task discovery, task belief encoding was inversely correlated with behavioral performance for shape category and was not correlated with response location. **a**, Follows Fig. 4e. Correlation between the neurally-estimated task belief (based on task classifier, see Methods) and shape category behavioral performance in C1 task for S1-C2-C1 task sequences. Behavioral performance is plotted relative to the S1 task. Therefore, lower numbers reflect behavior that is more aligned with the C1 task. Negative correlation reflects an increase in C1 task behavior when there is task belief is closer to the C1 task. **b**, Correlation between task belief encoding and color category encoding performance in C1 task for S1-C2-C1 task sequences. p ≤ 0.05, 0.01, and 0.001 for thin, medium, and thick lines, modified Mann-Kendall test, Benjamini-Hochberg correction for multiple comparisons. **c**, Follows Fig. 4k. Correlation between neurally-estimated task belief and decoded response direction during the C1 task for S1-C2-C1 task sequences. There was a weak correlation between stronger belief encoding and an increase in response direction encoding, but it was not significant.

## Methods

### Code availability

Custom MATLAB analysis functions are publicly available at https://github.com/buschman-lab/CompositionalTasks/.

### Subjects

Two adult male rhesus macaques (*Macaca mulatta*) participated in the experiment. Monkeys Si and Ch were between 8 and 11 years old and weighed approximately 12.7 and 10.7 kg, respectively. All experimental procedures were approved by the Princeton University Institutional Animal Care and Use Committee (protocol #3055) and were in accordance with the policies and procedures of the National Institutes of Health.

### Behavioral task

Each trial began with the monkeys fixating on a dot at the center of the screen. During a fixation period lasting 500ms–800ms, the monkeys were required to keep their gaze within a circle with a radius of 3.25 degrees of visual angle around the fixation dot. Following the fixation period, the stimulus and all four response locations were simultaneously displayed.

Stimuli were morphs consisting of both a color and shape (Fig. 1a). The stimuli were rendered as three-dimensional models using POV-Ray and MATLAB (Mathworks) and displayed on a Dell U2413 LCD monitor positioned 58 cm from the animal. Stimuli were morphed along circular continua in both color and shape (i.e., drawn from a four-dimensional ‘Clifford’ torus; Figure 1b). Colors were drawn from a photometrically isoluminant circle in the CIELAB color space, connecting the red and green prototype colors. Shapes were created by circularly interpolating the parameters defining the lobes of the ‘bunny’ prototype to the parameters defining the corresponding lobes of the ‘tee’ prototype. The mathematical representation of the morph levels adhered to the equation 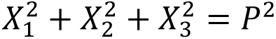 where *X* represents the parameter value in a feature dimension (e.g., L, a, b values in CIELAB color space). We chose the radius (*P*) to ensure sufficient visual discriminability between morph levels. The deviation of each morph level from prototypes (0% and 100%) was quantified using percentage, corresponding to positions on the circular space from −π *to* 0 and 0 *to* π. Morph levels were generated at eight levels: 0%, 30%, 50%, 70%, 100%, 70%, 50%, 30%, corresponding to 0, π/6, π/2, 5π/6, π, −5π/6, −π/2 and − π/6, respectively. 50% morph levels for one feature were only generated for prototypes of the other feature (i.e., 50% colors were only used with 0% or 100% shape stimuli). Stimuli were presented at fixation and had a diameter of 2.5 degrees of visual angle.

The monkeys indicated the color or shape category of the stimulus by saccading to one of the four response locations, positioned 6 degrees of visual angle from the fixation point at 45, 135, 225, and 315 degrees, relative to the vertical line. As the animals were performing the task, the reaction time was determined as the moment of leaving the fixation window relative to the time of stimulus onset.

Trials with a reaction time below 150ms were terminated, followed by a brief timeout (200ms). Correct responses were rewarded with juice, while incorrect responses resulted in short timeouts lasting from 1s for monkey Ch and 5s for monkey Si. After the trial finished, there was an inter-trial interval lasting 2–2.5 seconds before the next trial began.

The animals performed three category-response tasks (Fig. 1c). The Shape-Axis1 (S1) task required the monkeys to categorize the stimulus by its shape. For a stimulus with a shape that was closer to the ‘bunny’ prototype, the animals had to make a saccade to the upper-left (UL) location to get rewarded. For a stimulus with a shape closer to the ‘tee’ prototype, the animals had to make a saccade to the lower-right (LR) location to be rewarded. For ease of notation, we refer to the combination of the UL and LR target locations as ‘Axis 1’. The Color-Axis1 (C1) task required the monkeys to categorize the stimulus by its color. When a stimulus’ color was closer to ‘red’ the animals made an eye movement to the LR location and when closer to ‘green’ the animals made an eye movement to the UL location. Finally, in the Color-Axis2 (C2) task the animals again categorized stimuli based on their color but responded to the upper-right (UR) location for red stimuli and the lower-left (LL) location for green stimuli. Together, the UR and LL targets formed ‘Axis 2’. These set of three tasks were designed to be related to one another: the C1 and C2 tasks both required categorizing the color of the stimulus while the C1 and S1 tasks both required responding on Axis 1.

Animals were not explicitly cued as to which task was in effect. However, they did perform the same task for a block of trials allowing the animal to infer the task based on the combination of stimulus, response, and reward feedback. Tasks switched when the monkeys’ performance reached or exceeded 70% on the last 102 trials of Task S1 and Task C1 or the last 51 trials of Task C2. For Monkey Si, block switches occurred when their performance reached or exceeded the 70% threshold for all morphed and prototype stimuli in the relevant task dimension independently. For Monkey Ch, block switches occurred when their average performance at each morph level in the relevant task dimension exceeded the 70% threshold (e.g., average of all 30% morphs, average of all 70% morphs, and 0% and 100% prototypes in color dimension were all equal or greater than 70% accuracy in C1 and C2 tasks). In addition, on a subset of recording days, to prevent Monkey Ch from perseverating on one task for an extended period of time, the threshold was reduced to 65% over the last 75 trials for S1 and C1 tasks after the monkey had already done 200 or 300 trials on that block. When the animal hit the performance threshold, the task switched. This was indicated by a flashing yellow screen, a few drops of reward, and a delay of 50 secs.

To ensure even sampling of tasks despite the limited number of trials each day, the axis of response always changed following a block switch. During Axis 1 blocks, either S1 task or C1 task was pseudo-randomly selected, interleaved with C2 task blocks. Pseudo-random selection within Axis 1 blocks avoided three consecutive blocks of the same task, ensuring the animal performed at least one block of each task during each session.

Monkeys Si and Ch underwent training for 36 and 60 months, respectively. Electrophysiological recordings began when the monkeys consistently executed five or more blocks daily. Further details on the behavioral methods and results have been previously reported^1^.

### Congruent and incongruent stimuli

During the S1 and C1 tasks red-tee stimuli and green-bunny stimuli were ‘congruent’ as they required a saccade to the LR and UL locations, respectively, in both tasks. In contrast, stimuli in the red-bunny and green-tee portion of stimulus space were ‘incongruent’, as they required different responses during the two tasks (UL and LR in S1 task; LR and UL in C1 task, respectively). To ensure the animals performed the task well, 80% of the trials included incongruent stimuli. Note, as the C2 task was the only one to use Axis 2, there were no congruent or incongruent stimuli on those blocks.

### Analysis of behavioral data

Psychometric curves plot the fraction of trials in which the animals classified a stimulus with a specific morph level as being a member of the ‘green’ category for the C1 and C2 tasks or the ‘tee’ category for the S1 task. The average performance was calculated for each morph level during the last 102/102/51 trials of the S1/C1/C2 task. The fraction of responses for a given morph level of the task-relevant stimulus dimension was averaged across all morph levels of the task-irrelevant dimension (e.g., for the C1 task, the fraction of responses for 70% green stimuli were averaged across all shape morph levels). Since the stimulus space was circular, we averaged the behavioral response for the two stimuli at each morph level on each side of the circle (Fig. 1b). Psychometric curves were quantified by fitting the mean data points with a modified Gauss error function (erf):

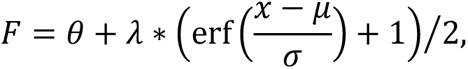

where erf is the error function, θ is a vertical bias parameter, λ is a squeeze factor, μ is a threshold, and *σ* is a slope parameter. Fitting was done in MATLAB with the Maximum likelihood method (fminsearch.m function).

To calculate performance during task discovery (Fig. 1f), we used a sliding window of 15 trials, stepped 1 trial, to calculate a running average of performance immediately after a switch or right before a switch. Performance was estimated using trials from all blocks of the same task, regardless of the identity of the stimulus.

### Surgical procedures and electrophysiological recordings

Monkeys were implanted with a titanium headpost to stabilize their head during recordings and two titanium chambers (19mm diameter) placed over frontal and parietal cortices that provided access to the brain. Chamber positions were determined using 3D model reconstruction of the skull and brain using structural MRI scans. We recorded neurons from lateral prefrontal cortex (LPFC, Brodmann area 46d and 46v), anterior inferior temporal cortex (aIT, area TEa), parietal cortex (PAR, Brodmann area 7), the frontal eye fields (FEF, Brodmann area 8) and the striatum (STR, caudate nucleus).

Two types of electrodes were used during recordings. To record from LPFC and FEF we used epoxy-coated tungsten single electrodes (FHC Inc., Bowdoin, ME). Pairs of single electrodes were placed in a custom-built grid with manual micromanipulators that lowered electrode pairs using a screw. This allowed us to record 20-30 neurons simultaneously from cortical areas near the surface. To record from deeper regions (PAR, aIT and STR) we used 16ch or 32ch Plexon V-Probes (Plexon Inc, Dallas, TX). These probes were lowered using the same custom-built grid through guide tubes. During lowering, we used both structural MRI scans and the characteristics of the electrophysiological signal to track the position of the electrode. A custom-made MATLAB GUI tracked electrode depth during the recording session and marked important landmarks until we found the brain region of interest.

Recordings were acute; up to 50 single electrodes and three V-Probes were inserted into the brain each day (in some recording sessions one additional V-Probe was also inserted in LPFC). Single electrodes and V-probes were lowered though the intact dura at the beginning of recording session and allowed to settle for 2-3 hours before recording, improving the stability of recordings. We did not simultaneously record from all five regions in all recording sessions. We began recording from LPFC and FEF for the first 5-10 days and then added PAR, STR, and aIT on successive days.

Broadband neural activity was recorded at 40 kHz using a 128-channel OmniPlex recording system (Plexon Inc., Dallas, TX). We performed 15 recording sessions with monkey Ch and 19 recording sessions with monkey Si. After all recordings were complete, we used electrical microstimulation in monkey Si, and structural MRI and microstimulation in monkey Ch, to identify FEF. Electrical stimulation was delivered as a train of anodal-leading biphasic pulses with a width of 400µs and an inter-pulse frequency of 330Hz. A site was identified as FEF when electrical microstimulation of ∼50µA evoked a stereotyped eye-movement on at least 50% of the stimulation attempts. In Monkey Ch, untested electrode locations were classified as FEF if they fell between two FEF sites (as confirmed with electrical stimulation) in a region that was confirmed as being FEF using MRI.

Eye position was recorded at 1 kHz using an Eyelink 1000 Plus eye-tracking system (SR Research, Kanata, ON). Sample onset was recorded using a photodiode attached to the screen. Eye position, photodiode signal, and behavioral events generated during the task, were all recorded alongside neural activity in the OmniPlex system.

### Signal preprocessing

Electrophysiological signals were filtered using a 4-pole high pass 300Hz Butterworth filter. To reduce common noise, the median of the signals recorded from all single electrodes in each chamber was subtracted from the activity of all single electrodes. For V-Probe recordings, we subtracted the median activity for all channels along the probe from each channel. To detect spike waveforms, a 4*n*_*n*_ threshold was used where *σ*_*n*_ is an estimate of standard deviation of noise distribution of signal *x* defined as:

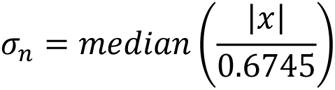

Time points at which the electrophysiological signal (*x*) crossed this threshold with a negative slope were identified as putative spiking events. Spike waveforms were saved (total length was 44 samples, 1.1ms, of which, 16 samples, 0.4ms, were pre-threshold). Repeated threshold crossing within 48 samples (1.2ms) were excluded. All waveforms recorded from a single channel were manually sorted into single units, multi-unit activity or noise using Plexon Offline Sorter (Plexon Inc., Dallas, TX). Units that were partially detected during a recording session were also excluded. All analyses reported in this manuscript were performed on single units.

For all reported electrophysiology analysis, saccade time was calculated as the moment at which the instantaneous eye speed exceeded a threshold of 720 degrees of visual angle per second. Instantaneous eye speed was calculated as 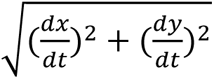, where *x* and *y* were position of the eye on the monitor at time *t*.

### Statistics and Reproducibility

Independent experiments were performed on two monkeys and the data were combined for subsequent analyses. As detailed below, statistical tests were corrected for multiple comparisons using two-tailed cluster correction unless stated otherwise. Unless otherwise noted, nonparametric tests were performed using 1,000 iterations; therefore, exact *p* values are specified when *p* > 0.001. Unless stated otherwise, all data were smoothed with a 150ms boxcar.

All analyses were performed in MATLAB 2021b (Mathworks).

### Using cluster mass to correct for multiple comparisons

To assess the significance of observed clusters in the time series data, we employed a two-tailed cluster mass correction method^2^. This approach is particularly useful when dealing with multiple comparisons and helps identify clusters of contiguous time points that exhibit statistically significant deviations from the null distribution.

We first generated a null distribution (N*ullDist*) by shuffling the observed data, breaking the relationship between the neural signal and the task parameter of interest (details of how data was shuffled are included with each test below). The observed data was also included as a ‘shuffle’ in the null distribution and the z-score of null distribution for each time point was calculated. To define significant moments in time, we computed the upper and lower thresholds based on the null distribution. The thresholds were determined non-parametrically using percentiles. The resulting thresholds, denoted as ‘*p*_*t*ℎr*es*ℎ*oldUppe*r’ and ‘*p*_*t*ℎr*es*ℎ*oldLowe*r’, serve as critical values for identifying significant deviations in both tails.

Timepoints of significant signal were identified by finding the moments when the value in each shuffle within the null distribution or the observed data exceeded the computed thresholds. These timepoints were then clustered in time, such that contiguous values above the threshold were summed together. The sum was calculated on the z-transformed values of each time series. This resulted in a ‘mass’ of each contiguous cluster in the data. To correct for multiple comparisons, we took the maximum absolute value of the cluster mass across time for each shuffle. This resulted in a distribution of maximum cluster masses. Finally, the two-tailed p-value of each cluster in the observed data was determined by comparing its mass to the distribution of maximum cluster masses in the null distribution.

### Firing rate (FR) calculation

In all analyses, we estimated the firing rate of each neuron by averaging the number of action potentials within a 100 ms window, stepping the window every 10ms. Changing the width of the smoothing window did not qualitatively change our results. The time labels in the figures denote the trailing edge of this moving window (i.e., 0 ms would be a window from 0 ms to 100 ms).

### Generalized Linear Model (GLM)

To estimate how individual neurons represented task variables, we used a Generalized Linear Model (GLM) to relate the activity of each neuron (*y*) to the task variables at each moment of time. The full model was:

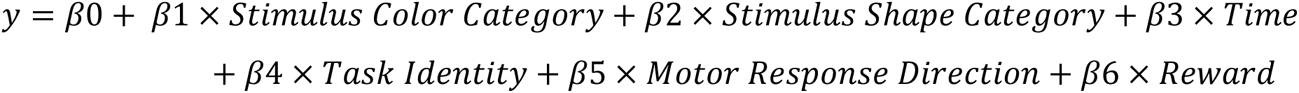

where *y* is the firing rate of each neuron, normalized by the max firing rate across all trials, and β0, β1, …, β6 are the regression coefficients corresponding to each predictor. Predictors were:

|  |  |
| --- | --- |
| Stimulus Color Category | Color category of the stimulus (categorical |
|  | variable) |
| Stimulus Shape Category | Shape category of the stimulus (categorical variable) |
| Time | The temporal progression within a recording session, normalized between 0 and 1 (continuous variable). |
| Task Identity | Identity of the task (categorical variable) |
| Motor Response Direction | Direction of the motor response (categorical variable; Upper-Left, Upper-Right, Lower-Left, or Lower-Right) |
| Reward | Whether a reward was received following the response (binary variable; 1 for reward, 0 for no reward) |

The GLM coefficients (β) were estimated using maximum likelihood estimation as implemented by MATLAB’s lassoglm.m function. Independent models were fit for each time point. To address potential overfitting, we applied Lasso (*L*^1^) regularization to the GLM weights with regularization coefficients (lambda) values of [0, 0.0003, 0.0006, 0.0009, 0.0015, 0.0025, 0.0041, 0.0067, 0.0111, 0.0183]. In a separate set of runs, we fit the GLM models to 80% of data and tested on the remaining 20%. The lambda value with maximum R^2^ on the withheld data was used when estimating the coefficient of partial determination (CPD).

### Coefficient of Partial Determination (CPD) Calculation

To quantify the unique contribution of each predictor to a neuron’s activity we used the Coefficient of Partial Determination (CPD). This metric quantifies the percent of explained variance that is lost when a specific factor is removed from the full model. The CPD was computed by initially fitting the GLM with all the factors (full model) as described above. Then, each factor was sequentially removed to create a set of reduced models, each of which were fit to the data. The CPD for each predictor *X*(*t*) was calculated as:

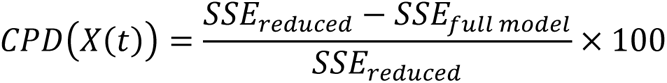

where *SSE*_r*educed*_ is the sum of square error due to predictor *X*, *SSE*_*full* *model*_is the sum of square error of the full model with all predictors^3^.

We used a permutation test to assess whether CPDs for each predictor, at each time-point, and for each neuron, was significantly larger than expected by chance. To compute a null distribution for CPD for each predictor, we generated 1000 permuted data sets by randomly permuting the predictor values relative to the neural activity and refitting the full and reduced models (as above). The likelihood of the observed CPD was then estimated by computing the proportion of permuted datasets that yielded a CPD greater than or equal to that of the observed dataset. To account for multiple comparisons across time-points, we used cluster correction (detailed above) to estimate corrected p-values for each predictor. Neurons that had at least one significant cluster (p-value < 0.05) were considered to significantly carry information about a task variable. To assess if an area had a significant number of neurons for a given task variable, we used a binomial test against the alpha-level 0.05.

For Fig. 1h-l, we compensated for a baseline CPD by subtracting the average permuted CPD from the observed CPD. For Fig. 1m, we averaged the CPD for each factor across all recorded neurons, subtracted the average baseline in –200-0ms since stimulus onset period and then normalized the resulting CPD curve to its maximum value. For Fig. S2, we averaged the CPD for task identity factor across all recorded neurons then normalized the resulting CPD curve to its maximum value. This highlighted the timing of different factors during the trial.

### Classifiers

To understand how task variables were represented in the neural population, we trained a set of binary classifiers to discriminate the vector of firing rates across the neural population for two different categories of task variables (depending on the variable of interest). Classifiers were trained with the logistic regression algorithm (as implemented in MATLAB, fitclinear.m function). In brief, the linear classifier relates *x* (vector of neural responses) to *y* (task labels, either +1 or –1) via a linear equation, with weights(*w*) and intercept(*b*):

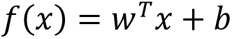

where *w* and *b* are optimized to minimize the logistic loss function: 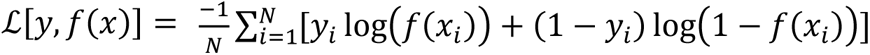, where N is the number of samples. Ridge (*L*^2^) regularization with a regularization coefficient λ = 1/60 was used to minimize over-fitting. The classifier was trained with the Broyden-Fletcher-Goldfarb-Shanno (BFGS) algorithm.

### Construction of pseudo-populations

All classifiers were trained on pseudo-population of responses constructed from neurons across all recording sessions. A separate pseudo-population was constructed for each classifier analysis based on the trial types of interest (described below). To be included in the pseudo-population, each neuron had to be recorded for a minimum number of trials (N_*t*r*ain*_ + N_*test*_) of each type of trial. The firing rate of neurons were concatenated to form a vector of neuron firing rates (the ‘pseudo-population’). Neural activity was aligned in time relative to either the sample onset or saccade onset, depending on the analysis.

To combine neurons across recording sessions into a single ‘trial’ of the pseudo-population, we drew trials for each neuron with matching experimental conditions (i.e., matched in terms of reward, color and shape morph level, task identity, and response direction). For instance, when constructing a pseudo-population for classifying the color category, the first ‘trial’ was constructed by concatenating the neural responses of neurons on trials that were rewarded, had a color morph level of 100, and a shape morph level of 30. If a neuron did not have a trial with an exactly matching stimulus, then a trial that matched the color category and reward would be randomly chosen.

### Cross-validation of classifiers

A separate logistic regression classifier was trained and tested on withheld trials for each time point. To estimate variability, the entire analysis was repeated 250 times, with each iteration involving a different partition of trials into training and testing sets. This new set of trials was randomly sampled with replacement (always ensuring test trials were separate from the train trials).

When testing the performance of a classifier trained on the same set of conditions (e.g., training and testing color categorization on the C2 task), the performance of the classifier was taken as the average performance across 10-fold cross-validation. However, as detailed in the main text, many of the classifiers were trained and tested across conditions (e.g., across different tasks or across trials during learning of the task). In this case, the test trials were randomly resampled 10 times for each training set, and the classifier’s performance was averaged across folds.

### Balancing classifier conditions

To avoid bias in the classifier, we balanced trial conditions to ensure the observed results were not due to other experimental factors. Conditions were balanced in two ways:

1. **Balanced congruency:** classifiers were trained on an equal number of congruent and incongruent trials, resulting in equal number of trials for four stimulus types “green-bunny”, “green-tee”, “red-bunny” and “red-tee”.
2. **Balanced reward**: classifiers were trained on an equal number of rewarded and unrewarded trials, balancing the stimulus identity of the relevant dimension on each side of the classifier.
3. **Balanced response direction**: classifiers were trained on an equal number of trials from each response direction on task specific axis (e.g. trials with response on Axis 1 for C1 task). This resulted in balanced response direction on each side of the classifier.

### Classifying the color category and shape category of stimuli

‘Color’ and ‘shape’ classifiers were trained to decode the stimulus color category or shape category from the vector of activity across the pseudo-population of neurons (Fig. 2a,b). A balanced number of congruent and incongruent stimuli were included in the training data to ensure color and shape information could be decoded independently. Classifiers were trained for each task independently.

Most classifiers were trained on correct trials alone. This maximized the number of trials included in the analysis and ensured the animals were engaged in the task. As many of our analyses were tested across tasks, this mitigates concerns that motor response information might confound stimulus category information. However, to ensure this was not the case, we controlled for motor response by balancing response direction in a separate set of analysis (Figs. S3a, S5a, and S7). While this significantly reduced the number of trials, and required us to include error trials, qualitatively similar results were often observed. Total number of LPFC, STR, aIT, PAR and FEF used for color category classification was 403, 110, 195, 54 and 116, respectively. Total number of LPFC, STR, aIT, PAR and FEF used for shape category classification was 480, 149, 239, 64 and 149, respectively.

### Classifying response direction

A separate set of ‘response’ classifiers were trained to decode the motor response from the vector of activity across the pseudo-population of neurons. Response direction was decoded within each axis (e.g., UL vs. LR for Axis 1). When training and testing on the same task (Fig. 2c), we only included trials that the animal responded on the correct axis (e.g., Axis 1 for the S1 and C1 tasks).

For testing whether response direction could be decoded across axes (Fig. S6a), we trained the classifier using trials from Task C1, where the animal responded on Axis 1, and then tested this classifier using trials from Task C2, where the animal responded on Axis 2, and vice versa. To control for stimulus-related information, we balanced rewarded and unrewarded trials for each condition and only included incongruent trials. This ensured that trials from the same stimulus category were present in both response locations. Similar results were seen when balancing for congruency and reward simultaneously, although the low number of incorrect trials for congruent trials resulted in a small set of neurons with the minimum number of train and test trials. Total number of LPFC, STR, aIT, PAR and FEF used for response direction classification was 403, 110, 195, 54 and 116, respectively. Due to limited number of trials, the total number of LPFC neurons for Fig. S6a right and Fig. S4c was 95 (only from animal Si).

### Using permutation tests to estimate the likelihood of classifier accuracy

We used permutation tests to estimate the likelihood the observed classifier accuracy occurred by chance. To create a null distribution of classifier performance, we randomly permuted the labels of training data 1000 times. Importantly, only the task-variable of interest was permuted – permutations were performed within the set of trials with the same identity of other (balanced) task variables. For example, if trials were balanced for reward, we shuffled labels within correct trials and within incorrect trials, separately. This ensured the shuffling only broke the relationship between the task-variable of interest and the activity of neurons. Shuffling of the labels was performed independently for each neuron before building the pseudo-population and training classifiers. Neurons that were recorded in the same session had identical shuffled labels. To stabilize the estimate of the classifier performance, the performance of the classifier on the observed and each instance of the permuted data were averaged over 10-folds and 20 to 50 novel iterations of each classifier. The null distribution was the combination of the permuted and observed values (total N=1001). The likelihood of the observed classifier performance was then estimated as its percentile within the null distribution. As detailed above, we used cluster correction to control for multiple comparisons across time. To depict significant time points within the task discovery period (highlighted as grey boxes in Figs. 4f, I, and 5a, b), we calculated the likelihood of classifier accuracy within consecutive windows of 50 trials (described below). Grey boxes were employed to denote significant time points, determined as those with a significance level of p ≤ 0.05 across all trial windows.

### Testing classification across tasks

To quantify whether the representation of color, shape, or response direction generalized across tasks (Fig. 2e,f,h,i), we trained classifiers on trials from one task and then tested the classifier on trials from another task. For example, to test cross-generalization of color information across the C1 and C2 tasks, a color classifier was trained on trials from the C2 task was tested on trials from the C1 task (Fig. 2f). To remove any bias due to differences in baseline firing rate across tasks, we subtracted the mean firing rate during each task from all trials of that task (at each time point). Similar results were observed when we did not subtract the mean firing rate.

### Cross-temporal classification

To measure how classifiers generalized across time, we trained classifiers to discriminate color category (Fig. 2e and Fig. S5c) or response direction (Fig. 2h) using 100ms time bins of FR data, sliding by 10ms. Classifiers trained on each time bin were then tested on all time bins of the test trials.

### Projection onto the encoding axis of classifier

To visualize the high-dimensional representation of task variables, we used the MATLAB predict.m function to project the firing rate response on test trials onto the one-dimensional encoding space defined by the vector orthogonal to the classification hyperplane. In other words, the projection measures the distance of the neural response vector to the classifier hyperplane. For example, to measure the encoding of each color category in the C1 task, we projected the trial-by-trial firing rate onto the axis orthogonal to the hyperplane of the color classifier trained during the C1 task.

### Quantifying the impact of task sequence on task discovery

The ‘task discovery’ period was defined as the first 125 trials after a switch in the task. The monkey’s performance increased during this period, as they learned which task was in effect (Fig. 1f). As detailed in the main manuscript, the animal’s behavioral performance depended on the sequence of tasks (Fig. 4a) and, so, we analyzed neural representations separately for two different sequences of tasks:

1. **C1-C2-C1 and S1-C2-S1 block sequence (‘same task transition’):** the task on Axis 1 (C1 or S1) repeated across blocks. As shown in the main text, monkeys tended to perform better during these task sequences, as if they were remembering the previous Axis 1 task.
2. **S1-C2-C1 and C1-C2-S1 block sequence (‘different task transition’):** the task on Axis 1 changed across blocks. As shown in the main text (Fig. 4a), monkeys tended to perform worse on these task sequences.

As we were interested in understanding how changes in neural representations corresponded to the animals’ behavioral performance, we divided the ‘different task transition’ sequences into two further categories: in ‘Low Initial Performance’ blocks, the animal’s behavioral performance during the first 25 trials of the block was less than 50% while on ‘High Initial Performance’ blocks performance was greater than 50%.

### Task belief representation during task discovery

To measure the animal’s internal representation of the task (i.e., their ‘belief’ about the task, as represented by the neural population), we trained a ‘task classifier’ to decode whether the current task was C1 or S1. The classifier was trained on neural data from the last 75 trials of each task block, when the animal’s performance was high (reflecting the fact that the animals were accurately estimating the task at the end of the block). The number of congruent and incongruent trials were balanced in the training dataset (32 trials: 4 trials for each of the four stimulus types, in each task). We included all C1 blocks regardless of their task sequence in training set. Only correct trials were used to train and test the classifier. To remove the effect of stimulus processing, the classifier was trained on neural activity from the fixation period (i.e., before stimulus onset^4^). As we were interested in measuring differences between tasks, we did not subtract the mean firing rate before training the classifier.

The task classifier was tested on trials from the beginning of blocks of the C1 task. Test trials were drawn from windows of 50 trials, slid every 5 trials, during learning (starting from trial 1-50 to trial 76-125). Overlapping test and train trials from the same task were removed. In contrast to training set, testing was done separately for S1-C2-C1 and C1-C2-C1 task sequences (Fig. 4b). As we were interested in focusing on the learning of the C1 task, we only used S1-C2-C1 task sequences with ‘low initial performance’ (see above). Classifiers were tested on pseudo-populations built from trials within each trial window, with a minimum of 4 test trials (1 trial for each of the four stimulus congruency types from Task C1). Neurons that did not include the required number of test trials for all trial windows were dropped. Note that, since we were using a small moving window of trials during task discovery, we had to trade-off the number of included neurons and the number of train/test trials for this analysis. In addition, although the number of correct trials increased as the animal discovered the task in effect, we kept a constant number of train and test trials in each of the sliding trial windows.

To quantify the animal’s belief above the current task, we measured the distance to the hyperplane of the C1/S1 task classifier (task belief encoding). For Fig. 4d, we averaged the performance of the classifier in pre-stimulus processing window (–400ms to 0ms).

### Correlation between task belief encoding and behavioral performance

To calculate the correlation between the animal’s task belief during task discovery and their behavioral performance (Fig. 4e), we used τ statistic in modified Mann-Kendall trend test (detailed below). This measurement was performed using data from each window of trials and the belief encoding, as estimated from the task classifier, on that same window of trials. As we are working with pseudo-populations of neurons, we estimated the behavioral performance for each window of trials during learning as the average of behavioral performance across all the trials in the window. The behavioral performance for each trial was taken as the average of the animal’s behavioral performance during the previous 10 trials. This yielded one average performance for each of the 16 trial windows. The task belief was measured for the same trials, using the average distance to the task classifier hyperplane, averaged over the time period from 400ms to 0ms before the onset of the stimulus (note that, as described above, all of these trials are withheld from the training data). Task belief was then taken as the average distance across all trials in a window of trials.

### Evolution of color category representation during task discovery

We were interested in quantifying how the shared color representation was engaged as the animal discovered the C1 task (Fig. 4f-g). To this end, we trained a classifier to categorize color using the last 75 trials of the C2 task, when the animal’s behavioral performance was high, and then tested it during learning of the C1 task. The classifier was trained on only correct trials and the training data was balanced for congruent and incongruent stimuli (16 trials: 4 trials for each of the four stimulus congruency types). As cross axis response decoding between the C1 and C2 tasks was weak (Fig. S6a), cross-task classifiers are capturing the representation of the color category that is shared between tasks.

Similar to the task classifier described above, the shared color classifier was tested using trials from the C1 task in a sliding window of 50 trials stepped 5 trials, in both the S1-C2-C1 and C1-C2-C1 sequences of tasks (Fig. 4b; as above, only low initial performance blocks were used for the S1-C2-C1 task sequences). We used 4 trials to test the classifier (1 trial for each of the four stimulus congruency types from Task C1). For Fig. 4g, we averaged the performance of the classifier in stimulus processing window (100ms to 300ms).

### Evolution of shape category representation during task discovery

We were interested in measuring the change in the representation of the stimulus’ shape category as the animal learned the C1 task (Fig. 4i). Our approach followed that of the color category representation described above, and so we only note differences here. We trained a classifier to categorize stimulus shape based on neural responses during the S1 task (limited to the last 75 trials of each block). To ensure the classifier was only responding to shape (and not motor response), we trained the classifier on a balanced set of correct (rewarded) and incorrect (unrewarded) trials (16 trials: 4 trials for each reward condition and for each shape category). The classifier was tested using trials from the C1 task (Fig. 4b). Note that we used the same C1 trials to quantify the representation of shared color category, shape category and task belief. For Fig. 4j, we averaged the performance of the classifier in stimulus processing window (100ms to 300ms). Total number of LPFC neurons included for S1-C2-C1 and C1-C2-C1 task sequences to quantify the representation of shared color category, shape category and task belief were 136 and 173, respectively.

### Evolution of response direction representation during task discovery

To measure the change in response direction representation as the monkey’s learned the C1 task, we trained a classifier to categorize the response direction based on the neural response during the S1 task (Fig. 4k; last 75 trials of the block). As above, we used a balanced number of correct and incorrect trials to control for information about the stimulus (12 trials: 3 trials for each reward condition and each response location). The classifier was then tested on correct trials from the C1 task during task learning (Fig. 4b; as above: sliding windows of 50 trials, stepped 5 trials, in S1-C2-C1 and C1-C2-C1; tested on 4 trials: 1 for each reward condition and each response location). For Fig. 4l, we averaged the performance of the classifier in response location processing window (200ms to 400ms). Total number of LPFC neurons included for S1-C2-C1 and C1-C2-C1 task sequences to quantify the representation of shared response direction were 120 and 155, respectively.

### Classifier statistical test to detect trend during task discovery period

We used a modified Mann Kendall test to quantify the statistical significance of trends in representations during task discovery (Figs. 4–5). The Mann-Kendall test is a non-parametric statistical test used to assess the presence of a monotonic trend in a time series data set^5,6^. This test is widely used in environmental and hydrological studies when quantifying trends in data. The null hypothesis is that there is no trend in the data (i.e., the data points are randomly ordered). The alternative hypothesis is that there is a monotonic trend, either increasing or decreasing. The test statistic, denoted as Tau (τ), represents the strength and direction of the trend. The original Mann-Kendall test assumes independence of data points. Here, we use a modified test that accounts for auto correlations within the data^7^. Specifically, we used Modified_MannKendall_test.m function, implemented in MATLAB, as previously described^8^. We set the alpha level for selecting significant autocorrelation lag values to 0.05 (alpha_ac=0.05). This ensured high sensitivity to autocorrelation in our analysis.

To measure significance of trends during task discovery, we first averaged the classifier accuracy across all repetitions for each time point (Figs. 4 and 5). We then calculated the Mann-Kendall p-value for each time point. To control for multiple comparisons over contiguous time points of a trial, we used Benjamini-Hochberg method (fdr_bh.m function, implemented in MATLAB, as previously described^9^).

### Transfer of information between subspaces

As detailed in the main text, we were interested in testing the hypothesis that the representation of the stimulus color in the shared color subspace predicted the response in the shared response subspace on a trial-by-trial basis (Figs. 3c-e and S6b-f). To this end, we correlated trial-by-trial variability in the strength of encoding of color and response along four different classifiers.

First, as detailed above, a shared color classifier was trained to decode color category from the C2 task and tested on trials from the C1 task. Training data were balanced for correct and incorrect trials (rewarded and unrewarded trials, 16 trials: 4 trials for each reward condition and each color category). Test trials were all correct trials (2 trials: 1 trial for each color category). Both train and test trials were drawn from the last 50 trials from the block (to ensure the animals were performing the task well). Total number of LPFC neurons included for this classifier was 63 (only from animal Si due to constraints on the number of trials).

Similarly, a second shared response classifier was trained on trials from the S1 task and then tested on the same set of test trials as the shared color classifier.

A third classifier was trained to decode the response direction on Axes 2, using a balance of correct and incorrect trials from the C2 task. This classifier was tested on the same set of test trials as the shared color classifier.

Finally, a fourth classifier was trained to decode the shared color representation but was now trained on correct trials from the C1 task (12 trials: 3 trials for each of the four stimulus congruency types) and tested on correct trials from C2 task (2 trials: 1 trial for each color category). Total number of LPFC neurons included for this classifier was 101 (only from animal Si, due to constraints on the number of trials).

To account for the arbitrary nature of positive and negative labels, we calculated the magnitude of encoding by flipping the encoding for negative labels. All four classifiers were trained on 2000 iterations of training set trials. Note, that the first three classifiers were tested on the same test trials, allowing for trial-by-trial correlations to be measured. Furthermore, all four classifiers were trained over time, allowing us to measure the cross-temporal correlation between any pairs of classifiers.

Altogether, these four classifiers allowed us to test three hypothesized correlations that reflect the transfer of task-relevant stimulus information into representations of behavioral responses. First, one might expect that, on any given trial before the start of saccade, the strength and direction of the shared color representation, as estimated by the distance to the hyperplane of the first classifier, should be correlated with the shared response, as estimated by the distance to the hyperplane of the second classifier. This correlation is seen in Fig. 3c.

Second, during the performance of the C1 task, one would expect that before the start of saccade the shared color representation to not be correlated with the response on the Axes 2 predicted by the C2 task. This is quantified by correlating the distance to the hyperplane of the first classifier and the distance to the hyperplane of the third classifier, as seen in Fig. 3d.

However, one would expect these representations to be correlated when the animal is performing the C2 task. This is quantified by correlating the distance to the hyperplane of the fourth classifier and the distance to the hyperplane of the third classifier, as seen in Fig. S6b.

### Distance along color and shape encoding axes during the discovery of the task

To understand how the geometry of the neural representation of the color and shape of the stimulus evolved during task discovery (Figure 5), we measured the distance between the two prototype colors (red and green) and the two prototype shapes (tee and bunny). Distance was measured along the encoding axis for each stimulus dimension (i.e., the axis that is orthogonal to the color and shape classifiers, described above). So, for each test trial, the distance along the color encoding axis was:

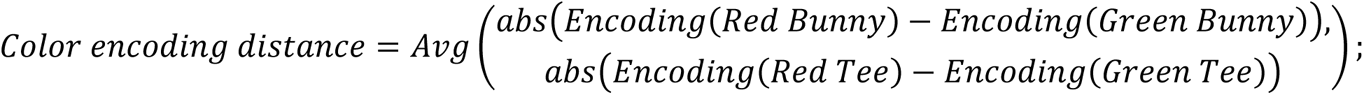

This approach allowed us to calculate the distance along red/green colors while controlling for differences across shapes. The color encoding distance was estimated for 250 iterations of the classifiers, allowing us to estimate the mean and standard error of the distance. A similar process was followed for estimating the shape encoding distance.

To calculate the color and shape encoding distance during task discovery (Fig. 5a, 5b), we calculated the distance above in the sliding window of 50 trials after the switch into the C1 task.

### Compression index (CPI)

Compression index (CPI) was defined as the log of the ratio of average “Color encoding distance” and average “Shape encoding distance” described above:

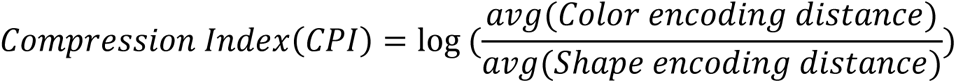

To calculate CPI for each task (Fig. 5c), we computed shape and color encoding distance using all trials of a given task. To measure the shape distance for all three tasks, we trained a classifier to categorize stimulus shape based on neural responses during the S1 task (limited to the last 75 trials of each block). To ensure the classifier was only responding to shape (and not motor response), we trained the classifier on a balanced set of correct (rewarded) and incorrect (unrewarded) trials (20 trials: 5 trials for each reward condition and for each shape category). Shape encoding for each task was calculated using 4 trials to test the classifier (1 trial for each of the four stimulus congruency types).

To measure color distance in S1/C2/C1 task, we trained color category classifier on only correct trials of C1/C1/C2 task balanced for congruent and incongruent stimuli (20 trials: 5 trials for each of the four stimulus congruency types). Color encoding for each task was calculated using 4 trials of that task to test the classifier (1 trial for each of the four stimulus congruency types).

To track changes in the relative strength of color and shape information during task discovery, we calculated CPI in a sliding window of 50 trials after the animal’s switched into the C1 task (during S1-C2-C1 sequences of tasks, with low initial performance). Note, since we controlled for motor response when calculating shape and color distance, the CPI is not affected by motor response information^10^.

### Correlation between task belief encoding and compression index

To correlate the CPI with task belief encoding, we used the same set of C1 test trials to calculate both CPI and estimate the task identity using the task classifier described above. CPI values in 100ms-300ms after stimulus onset and belief encoding values in 400ms to 0ms before stimulus onset were averaged for all trials in each trial window and the Mann-Kendall correlation between the resulting two vectors was calculated (Fig. 5e).

## References

1. Sakai, K. Task Set and Prefrontal Cortex. Annual Review of Neuroscience 31, 219–245 (2008).

2. Yang, G. R., Joglekar, M. R., Song, H. F., Newsome, W. T. & Wang, X.-J. Task representations in neural networks trained to perform many cognitive tasks. Nat Neurosci 22, 297–306 (2019).

3. Driscoll, L., Shenoy, K. & Sussillo, D. Flexible Multitask Computation in Recurrent Networks Utilizes Shared Dynamical Motifs. http://biorxiv.org/lookup/doi/10.1101/2022.08.15.503870 (2022) doi:10.1101/2022.08.15.503870.

4. Goudar, V., Peysakhovich, B., Freedman, D. J., Buffalo, E. A. & Wang, X.-J. Schema formation in a neural population subspace underlies learning-to-learn in flexible sensorimotor problem-solving. Nat Neurosci 1–12 (2023) doi:10.1038/s41593-023-01293-9.

5. Epstein, R., Kirshnit, C. E., Lanza, R. P. & Rubin, L. C. ‘Insight’ in the pigeon: antecedents and determinants of an intelligent performance. Nature 308, 61–62 (1984).

6. Ito, T. et al. Compositional generalization through abstract representations in human and artificial neural networks. Preprint at http://arxiv.org/abs/2209.07431 (2022).

7. Makino, H. Arithmetic value representation for hierarchical behavior composition. Nat Neurosci 1–10 (2022) doi:10.1038/s41593-022-01211-5.

8. Reverberi, C., Görgen, K. & Haynes, J.-D. Compositionality of Rule Representations in Human Prefrontal Cortex. Cerebral Cortex 22, 1237–1246 (2012).

9. Tenenbaum, J. B., Kemp, C., Griffiths, T. L. & Goodman, N. D. How to Grow a Mind: Statistics, Structure, and Abstraction. Science 331, 1279–1285 (2011).

10. Frankland, S. M. & Greene, J. D. Concepts and Compositionality: In Search of the Brain’s Language of Thought. Annu. Rev. Psychol. 71, 273–303 (2020).

11. Botvinick, M. et al. Reinforcement Learning, Fast and Slow. Trends in Cognitive Sciences 23, 408–422 (2019).

12. Lake, B. M., Ullman, T. D., Tenenbaum, J. B. & Gershman, S. J. Building Machines That Learn and Think Like People. Preprint at 10.48550/arXiv.1604.00289 (2016).

13. Lake, B. M. & Baroni, M. Human-like systematic generalization through a meta-learning neural network. Nature 1–7 (2023) doi:10.1038/s41586-023-06668-3.

14. Lin, B., Bouneffouf, D. & Rish, I. A Survey on Compositional Generalization in Applications. Preprint at http://arxiv.org/abs/2302.01067 (2023).

15. Marcus, G. The Next Decade in AI: Four Steps Towards Robust Artificial Intelligence. arXiv.org https://arxiv.org/abs/2002.06177v3 (2020).

16. Bouchacourt, F., Tafazoli, S., Mattar, M. G., Buschman, T. J. & Daw, N. D. Fast rule switching and slow rule updating in a perceptual categorization task. eLife 11, e82531 (2022).

17. Siegel, M., Buschman, T. J. & Miller, E. K. Cortical information flow during flexible sensorimotor decisions. Science 348, 1352–1355 (2015).

18. Mante, V., Sussillo, D., Shenoy, K. V. & Newsome, W. T. Context-dependent computation by recurrent dynamics in prefrontal cortex. Nature 503, 78–84 (2013).

19. Musall, S., Kaufman, M. T., Juavinett, A. L., Gluf, S. & Churchland, A. K. Single-trial neural dynamics are dominated by richly varied movements. Nat. Neurosci. 22, 1677–1686 (2019).

20. Talluri, B. C. et al. Activity in primate visual cortex is minimally driven by spontaneous movements. Nat. Neurosci. 26, 1953–1959 (2023).

21. MacDowell, C. J., Libby, A., Jahn, C. I., Tafazoli, S. & Buschman, T. J. Multiplexed Subspaces Route Neural Activity Across Brain-wide Networks. 2023.02.08.527772 Preprint at 10.1101/2023.02.08.527772 (2023).

22. Wallis, J. D., Anderson, K. C. & Miller, E. K. Single neurons in prefrontal cortex encode abstract rules. Nature 411, 953–956 (2001).

23. Barbosa, J. et al. Early selection of task-relevant features through population gating. Nat Commun 14, 6837 (2023).

24. Flesch, T., Juechems, K., Dumbalska, T., Saxe, A. & Summerfield, C. Orthogonal representations for robust context-dependent task performance in brains and neural networks. Neuron 110, 1258–1270.e11 (2022).

25. Langdon, C. & Engel, T. A. Latent circuit inference from heterogeneous neural responses during cognitive tasks. 2022.01.23.477431 Preprint at 10.1101/2022.01.23.477431 (2022).

26. Miller, E. K. & Cohen, J. D. An Integrative Theory of Prefrontal Cortex Function. Annual Review of Neuroscience 24, 167–202 (2001).

27. Takagi, Y., Hunt, L. T., Woolrich, M. W., Behrens, T. E. & Klein-Flügge, M. C. Adapting non-invasive human recordings along multiple task-axes shows unfolding of spontaneous and over-trained choice. eLife 10, e60988 (2021).

28. Mack, M. L., Preston, A. R. & Love, B. C. Ventromedial prefrontal cortex compression during concept learning. Nature Communications 11, 1–11 (2020).

29. Wójcik, M. J., et al. Learning Shapes Neural Geometry in the Prefrontal Cortex. 10.1101/2023.04.24.538054 (2023) doi:10.1101/2023.04.24.538054.

30. Xue, C., Kramer, L. E. & Cohen, M. R. Dynamic task-belief is an integral part of decision-making. Neuron (2022) doi:10.1016/j.neuron.2022.05.010.

31. Rigotti, M. et al. The importance of mixed selectivity in complex cognitive tasks. Nature 497, 585– 590 (2013).

32. Rouzitalab, A., Boulay, C. B., Park, J., Martinez-Trujillo, J. C. & Sachs, A. J. Ensembles code for associative learning in the primate lateral prefrontal cortex. Cell Reports 42, (2023).

33. Barak, O., Rigotti, M. & Fusi, S. The Sparseness of Mixed Selectivity Neurons Controls the Generalization–Discrimination Trade-Off. J. Neurosci. 33, 3844–3856 (2013).

34. Gurnani, H. & Cayco Gajic, N. A. Signatures of task learning in neural representations. Current Opinion in Neurobiology 83, 102759 (2023).

35. Turner, E. & Barak, O. The Simplicity Bias in Multi-Task RNNs: Shared Attractors, Reuse of Dynamics, and Geometric Representation. Adv. Neural Inf. Process. Syst. 36, 25495–25507 (2023).

36. Muhle-Karbe, P. S. et al. Goal-seeking compresses neural codes for space in the human hippocampus and orbitofrontal cortex. Neuron 111, 3885–3899.e6 (2023).

37. Bernardi, S. et al. The Geometry of Abstraction in the Hippocampus and Prefrontal Cortex. Cell 183, 954–967.e21 (2020).

38. Panichello, M. F. & Buschman, T. J. Shared mechanisms underlie the control of working memory and attention. Nature 592, 601–605 (2021).

39. Johnston, W. J. & Fusi, S. Abstract representations emerge naturally in neural networks trained to perform multiple tasks. Nat Commun 14, 1040 (2023).

40. Whittington, J. C. R. et al. The Tolman-Eichenbaum Machine: Unifying Space and Relational Memory through Generalization in the Hippocampal Formation. Cell 183, 1249–1263.e23 (2020).

41. Baars, B. J. Global workspace theory of consciousness: toward a cognitive neuroscience of human experience. in Progress in Brain Research vol. 150 45–53 (Elsevier, 2005).

42. Kaufman, M. T., Churchland, M. M., Ryu, S. I. & Shenoy, K. V. Cortical activity in the null space: permitting preparation without movement. Nat Neurosci 17, 440–448 (2014).

43. Libby, A. & Buschman, T. J. Rotational dynamics reduce interference between sensory and memory representations. Nat Neurosci 24, 715–726 (2021).

44. Hummos, A. Thalamus: a brain-inspired algorithm for biologically-plausible continual learning and disentangled representations. Preprint at http://arxiv.org/abs/2205.11713 (2023).

45. Jahn, C. I., Markov, N. T., Morea, B., Ebitz, R. B. & Buschman, T. J. Learning attentional templates for value-based decision-making. Cell (2024).

46. Niv, Y. Reinforcement learning in the brain. Journal of Mathematical Psychology 53, 139–154 (2009).

47. Singh, D., Norman, K. A. & Schapiro, A. C. A model of autonomous interactions between hippocampus and neocortex driving sleep-dependent memory consolidation. Proc. Natl. Acad. Sci. U.S.A. 119, e2123432119 (2022).

## Supplemental References

1. Bouchacourt, F., Tafazoli, S., Mattar, M. G., Buschman, T. J. & Daw, N. D. Fast rule switching and slow rule updating in a perceptual categorization task. eLife 11, e82531 (2022).

2. Maris, E. & Oostenveld, R. Nonparametric statistical testing of EEG– and MEG-data. Journal of Neuroscience Methods 164, 177–190 (2007).

3. Chiang, F.-K. & Wallis, J. D. Neuronal Encoding in Prefrontal Cortex during Hierarchical Reinforcement Learning. 30, 12.

4. Wallis, J. D., Anderson, K. C. & Miller, E. K. Single neurons in prefrontal cortex encode abstract rules. Nature 411, 953–956 (2001).

5. Kendall, M. G. Rank Correlation Methods. (Griffin, Oxford, England, 1948).

6. Mann, H. B. Nonparametric Tests Against Trend. Econometrica 13, 245–259 (1945).

7. Hamed, K. H. & Ramachandra Rao, A. A modified Mann-Kendall trend test for autocorrelated data. Journal of Hydrology 204, 182–196 (1998).

8. Aalok, A (2024). Modified-MannKendall-Test (https://github.com/atharvaaalok/Modified-MannKendall-Test/releases/tag/v2.0), GitHub. January, 2024.

9. Groppe, D (2024). fdr_bh (https://www.mathworks.com/matlabcentral/fileexchange/27418-fdr_bh), MATLAB Central File Exchange. January, 2024.

10. Flesch, T. et al. Are task representations gated in macaque prefrontal cortex? Preprint at http://arxiv.org/abs/2306.16733 (2023).

